# Adeno-to-squamous transition drives resistance to KRAS inhibition in *LKB1* mutant lung cancer

**DOI:** 10.1101/2023.09.07.556567

**Authors:** Xinyuan Tong, Ayushi S. Patel, Eejung Kim, Hongjun Li, Yueqing Chen, Shuai Li, Shengwu Liu, Julien Dilly, Kevin S. Kapner, Yun Xue, Laura Hover, Suman Mukhopadhyay, Fiona Sherman, Khrystyna Mynzdar, Yijun Gao, Fei Li, Fuming Li, Zhaoyuan Fang, Yujuan Jin, Juntao Gao, Minglei Shi, Luonan Chen, Yang Chen, Thian Kheoh, Wenjing Yang, Itai Yanai, Andre L. Moriera, Vamsidhar Velcheti, Benjamin G. Neel, Liang Hu, James G. Christensen, Peter Olson, Dong Gao, Michael Q. Zhang, Andrew J. Aguirre, Kwok-Kin Wong, Hongbin Ji

**Affiliations:** State Key Laboratory of Cell Biology, Shanghai Institute of Biochemistry and Cell Biology, Center for Excellence in Molecular Cell Science, Chinese Academy of Sciences, Shanghai 200031, China; Laura and Isaac Perlmutter Cancer Center, New York University Langone Health, New York, NY 10016, United States; Department of Medical Oncology, Dana-Farber Cancer Institute, Boston, MA 02215, United States; Broad Institute of MIT and Harvard, Cambridge, MA 02142, United States; Department of Medicine, Brigham and Women’s Hospital and Harvard Medical School, Boston, MA 02115, United States; MOE Key Laboratory of Bioinformatics, Bioinformatics Division and Center for Synthetic and Systems Biology, BNRist, Department of Automation, Tsinghua University, Beijing 100084, China; University of Chinese Academy of Sciences, Beijing 100049, China; School of Life Science, Hangzhou Institute for Advanced Study, University of Chinese Academy of Sciences, Hangzhou 310024, China; Monoceros Biosystems, LLC, San Diego, CA 92129, United States; State Key Laboratory of Oncology in South China, Collaborative Innovation Center for Cancer Medicine, Sun Yat-Sen University Cancer Center, Guangzhou, Guangdong 510060, China; Department of Pathology, School of Basic Medical Sciences, Fudan University, Shanghai 200032, China; Shanghai Key Laboratory of Metabolic Remodeling and Health, Institute of Metabolism and Integrative Biology, Fudan University, Shanghai 200438, China; University of Edinburgh Institute, Zhejiang University, Haining 314400, China; MOE Key Laboratory of Bioinformatics, Bioinformatics Division and Center for Synthetic and Systems Biology, BNRist, School of Medicine, Tsinghua University, Beijing 100084, China; School of Life Science and Technology, Shanghai Tech University, Shanghai 200120, China; Key Laboratory of Systems Biology, Hangzhou Institute for Advanced Study, University of Chinese Academy of Sciences, Chinese Academy of Sciences, Hangzhou 310024, China; Center for Excellence in Animal Evolution and Genetics, Chinese Academy of Sciences, Kunming 650223, China; State Key Laboratory of Medical Molecular Biology, Department of Molecular Biology and Biochemistry, Institute of Basic Medical Sciences, Chinese Academy of Medical Sciences, School of Basic Medicine, Peking Union Medical College, Beijing 100049, China; Mirati Therapeutics, San Diego, CA 92121, United States; Institute of Systems Genetics, New York University Langone Health, New York, NY 10016, United States; Department of Biological Sciences, Center for Systems Biology, The University of Texas, Richardson, TX 75080, United States

**Keywords:** Adeno-to-squamous transition (AST), KRAS inhibitor, LKB1, Organoid, KRT6A

## Abstract

KRAS^G12C^ inhibitors including adagrasib and sortorasib have shown clinical promise in targeting *KRAS^G12C^*-mutated lung cancers, however, most patients eventually develop drug resistance. In lung adenocarcinoma patients with co-occurring *KRAS^G12C^* and *STK11*/*LKB1* mutations, we found a high squamous gene signature at baseline significantly correlated with poor adagrasib response. Through integrative studies of *Lkb1*-deficient *KRAS^G12C^*and *Kras^G12D^* lung cancer mouse models and/or organoids treated with KRAS inhibitors, we found tumor cells invoked a lineage plasticity program: adeno-to-squamous transition (AST) that mediated resistance to KRAS inhibition. Transcriptomic and epigenomic analyses revealed ΔNp63 drives AST and modulates response to KRAS inhibition. We identified an intermediate high-plasticity cell state with distinct gene expression program marked by *Krt6a* upregulation. Notably, higher *KRT6A* expression at baseline correlated with shorter overall survival in *KRAS*-mutant patients receiving adagrasib. These data support the role of AST in KRAS inhibitor resistance and provide predictive biomarker for KRAS-targeted therapies in lung cancer.

## Introduction

*KRAS* is one of the most frequently mutated genes in lung ADC (Riely et al., 2009; The Cancer Genome Atlas Research Network, 2014) and is frequently mutated at codons 12, 13 and 61 (Karoulia et al., 2017). *KRAS* mutations have been shown to act as oncogenic drivers by mediating the activation of the downstream signaling, including the rapidly accelerated fibrosarcoma (RAF), mitogen-activated protein kinase kinase (MEK) and extracellular regulated protein kinases (ERK) pathway. While *KRAS* has long been considered a challenging therapeutic target (Cox et al., 2014), recent development of covalent inhibitors of the inactive, GDP-bound form of the KRAS^G12C^ mutant protein has transformed therapeutic strategies for tumors harboring this mutation (Ryan and Corcoran, 2018). Two KRAS^G12C^ inhibitors – sotorasib (AMG510) and adagrasib (MRTX849) – have demonstrated clinical efficacy in *KRAS^G12C^*-mutant lung cancer and are now approved for therapeutic use in the second-line or beyond setting (Canon et al., 2019; Hallin et al., 2020; Hong et al., 2020; Janes et al., 2018; Kim et al., 2020; Skoulidis et al., 2021). Additionally, early signs of efficacy have been observed for these KRAS^G12C^ inhibitors in colorectal and pancreatic cancers (Strickler et al., 2023; Yaeger et al., 2023). Moreover, inhibitors against other *KRAS* mutations including *KRAS*^G12D^ are being developed (e.g. MRTX1133) (Hallin et al., 2022; Kemp et al., 2023; Wang et al., 2022). While these therapeutic strategies have been effective in treating *KRAS^G12C^*-mutant lung cancer (Jänne et al., 2022; Skoulidis et al., 2021), primary and acquired resistance to therapy are common, and it is critical to define the mechanisms of resistance to KRAS inhibition to fuel effective combination therapy strategies.

A number of studies have reported genetic and non-genetic mechanisms of resistance to KRAS^G12C^ inhibitors (Blaquier et al., 2021). We and others have found unique mutations in *KRAS* occurring both in *cis* and in *trans KRAS* allele (Awad et al., 2021; Koga et al., 2021). These include novel mutations within the adagrasib-binding pocket of *KRAS.* Moreover, mutations in multiple effectors of RAS/MAPK pathway including *NRAS*, *BRAF* and *MAP2K1* have been observed that all result in RAS pathway reactivation (Fedele et al., 2021; Tanaka et al., 2021; Xue et al., 2020; Zhao et al., 2021). Non-genetic mechanisms of resistance have also been observed, including those resulting in epithelial-to-mesenchymal transition (EMT) (Adachi et al., 2020), alteration of cell cycle (Hallin et al., 2020) and rewiring of metabolic pathways, such as fatty acid metabolism (Tsai et al., 2022). We recently described mechanisms of acquired resistance to adagrasib monotherapy and showed that histological transition from ADC to squamous cell carcinoma (SCC) might serve as a mechanism of resistance (Awad et al., 2021). In this study, two out of nine lung ADC patients with repeat tumor biopsies at the time of acquired resistance showed evidence of SCC histology without an evidence of a concurrent genomic alteration that would explain resistance (Awad et al., 2021). While these data indicated the potential link between AST and KRAS^G12C^ inhibitor resistance, additional preclinical and clinical data are needed to define the causal role of AST in mediating resistance.

Previous studies have noted AST as an emerging mechanism of resistance to other targeted therapies (Chen et al., 2019; Cooper et al., 2022; Passaro et al., 2021; Schoenfeld et al., 2020). For instance, AST is observed in 8% (5/62) of lung ADC patients relapsed from osimertinib treatment, a third-generation EGFR tyrosine kinase inhibitor (TKI) (Schoenfeld et al., 2020). While AKT pathway activation and aberrant activity of the MYC and SOX families of transcription factors have been implicated as being differentially activated between pre- and post-TKI-treatment (Quintanal-Villalonga et al., 2021), the mechanisms underlying AST as a consequence of KRAS inhibition still remain to be elucidated.

Genetically engineered mouse models (GEMMs) based on two common *KRAS* mutations, including *G12C* and *G12D* substitutions (Cerami et al., 2012; Cook et al., 2021; Riely et al., 2009), e.g., *KRAS ^LSL-G12C/+^* (Li et al., 2018) or *Kras ^LSL-G12D/+^* (Jackson et al., 2001), have been established to recapitulate human lung ADC. *STK11* (serine-threonine kinase 11)/*LKB1* is often concurrently mutated with *KRAS* in 15%-30% of human lung ADC (Sanchez-Cespedes et al., 2002), and *Lkb1* knockout in mice promotes lung cancer progression (Ji et al., 2007; Mahoney et al., 2009). LKB1 is a central regulator of chromatin accessibility, causing reduced accessibility at genomic regions with NKX2 motifs and increased accessibility at genomic regions with SOX motifs (Pierce et al., 2021). Interestingly, *Lkb1* deficiency induces squamous transition of *Kras^G12D^*-mutated lung ADC (Han et al., 2014; Zhang et al., 2017). This process was associated with progressively increased expression of squamous markers including ΔNp63, SOX2 and KRT5/14, and gradually decreased expression of NKX2-1 (Fang et al., 2023; Gao et al., 2014b; Tang et al., 2023).

In this study, we demonstrated that higher baseline expression of SCC genes was correlated with a shorter treatment duration with adagrasib in *KRAS^G12C^*;*LKB1* mutated non-small cell lung cancer (NSCLC) patients enrolled in the KRYSTAL-1 clinical trial. Furthermore, we showed that *KRAS^G12C^*;*Lkb1* mutant mouse models of lung ADC specifically invoked cell state plasticity when treated with adagrasib and exhibited squamous transition. We further established the role of AST in mediating resistance to KRAS inhibition in organoid models of *Kras^G12D^* or *KRAS^G12C^* with *Lkb1* loss. We identified two subtypes of *Lkb1*-mutant ADC in organoid models, including those with high or low squamous gene expression features with differential responses to KRAS inhibitors. Mechanistically, we identified ΔNp63 as the regulator of squamous transition and response to KRAS inhibition. Finally, we nominate an AST plasticity-related gene, *KRT6A,* to be a potential biomarker of a poor response to KRAS^G12C^ inhibitors in lung cancer.

## Results

### SCC signature correlates with adagrasib resistance in *KRAS^G12C^*;*LKB1* mutant NSCLC

Our clinical trial data have previously implicated a potential involvement of squamous transition in acquired resistance to adagrasib treatment in NSCLC patients (Awad et al., 2021). To interrogate the relationship between SCC-associated gene expression with the drug response in *KRAS^G12C^*-mutant lung ADC, we further analyzed the transcriptomic and clinical data from 116 lung cancer patients enrolled in KRYSTAL-1 trial, a phase I/II study of adagrasib in *KRAS^G12C^-*mutated cancers (Jänne et al., 2022). Available clinical data included treatment duration, progression free survival (PFS) and overall survival (OS) with adagrasib monotherapy. A total of 68 patients had available whole transcriptome profiling data from a pre-treatment biopsy (Methods). We scored each patient’s tumor for ADC and SCC related genes as well as for the SCC gene signature (Inamura et al., 2005), and evaluated correlation with clinical variables (Table S1-S3).

Interestingly, we observed that patients with tumors that had higher expression of SCC related genes *KRT14*, *KRT6A* and *KRT6B* generally spent a shorter time on adagrasib treatment, while patients with tumors with higher expression of the ADC-related genes *NAPSA*, *NKX2-1*, *SFTPB* trended towards longer adagrasib treatment duration (Table S3). Notably, we found that patients with tumors expressing higher enrichment of the SCC signature at baseline had a statistically significant shorter treatment duration with adagrasib (Figure 1A). We further investigated the association of the SCC signature enrichment with clinical outcomes on adagrasib therapy. Stratifying patients by the upper and lower quartile of SCC signature enrichment, i.e. high- and low-SCC scores, we observed a significantly shorter duration of treatment in the SCC-high cohort (Figure S1A).

**Figure 1:**
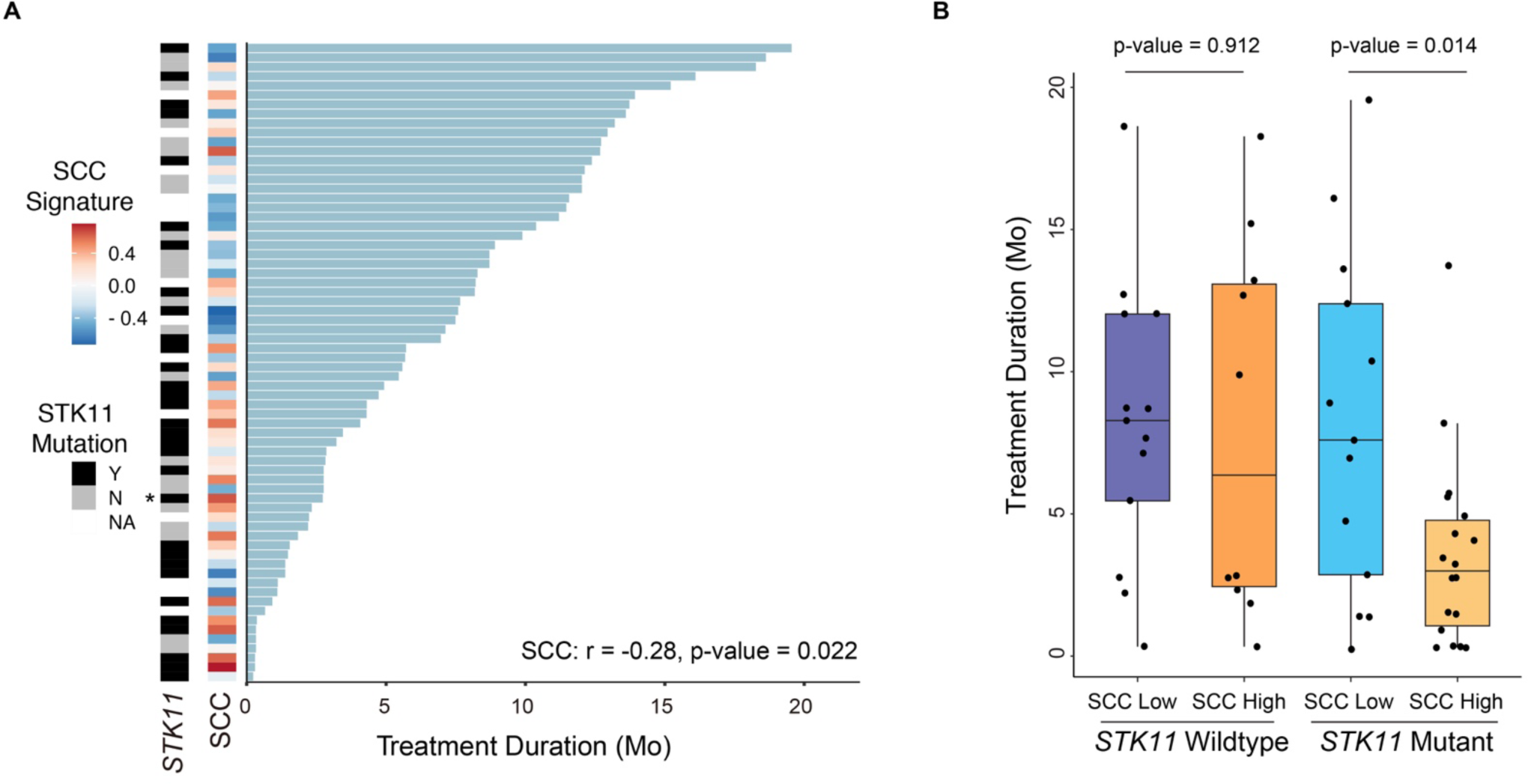
SCC program enrichment in pre-treatment biopsies associates with adverse outcomes to adagrasib monotherapy in lung cancer. **A.** Swimmer’s Plot showing correlation between SCC signature for each patient and time on treatment. Both SCC signature and STK11 mutation status are shown on the left. All patients are with ADC except for one SCC patient indicated with *. **B.** Boxplot showing the range of treatment duration stratified by *STK11*/*LKB1* mutation status and SCC signature scores above the median (high) and below the median (low). Two-tailed t-test was used for statistical analyses.

Given that *STK11*/*LKB1* mutations frequently co-occur with *KRAS^G12C^* mutations in lung ADC (Figure S1B) and prior studies have suggested a role for LKB1 in regulating cellular plasticity (Han et al., 2014; Pierce et al., 2021; Zhang et al., 2017), we sought to investigate the relationship between SCC signature and *STK11/LKB1* mutation status. We found that patients harboring *STK11-*mutant tumors that were enriched for the SCC signature experienced a significantly shorter treatment duration with adagrasib compared to those with low scores of the SCC signature; a similar association was not observed within the *STK11/LKB1* wildtype subgroup (Figures 1B and S1C). Moreover, in the two lung cancer patients who developed acquired resistance to adagrasib in our previous study (Awad et al., 2021), one patient had *STK11/LKB1* mutation, further supporting the potential role of *STK11/LKB1* in AST. These data suggest that patients harboring *STK11/LKB1-*mutant lung ADC with high baseline expression of the SCC signature may have worse outcomes with adagrasib monotherapy compared to those with low baseline expression of the SCC signature.

### Adagrasib-resistant mouse *KRAS^G12C^*/*Lkb1*-loss tumors harbor SCC features

Based on the identification of *KRAS*;*LKB1* mutant lung ADC tumors with high expression of SCC markers that experience with shorter duration on adagrasib therapy, we hypothesized that this subset of *KRAS*;*LKB1* mutated patients has a higher degree of cell state plasticity that renders them susceptible to therapy-induced squamous transition. To investigate this as a mechanism of resistance to KRAS^G12C^ inhibition, we employed two lung cancer GEMMs, *KRAS^LSL-G12C/+^/Trp53^flox/flox^* (K_C_P) and *KRAS^LSL-G12C/+^/Lkb1^flox/flox^* (K_C_L) models – representing the *TP53* and *STK11/LKB1* mutant genotypes, two of the most frequently co-mutated tumor suppressors in human lung ADC in conjunction with *KRAS* (Figure S1B). We induced tumorigenesis in both K_C_P and K_C_L models by intra-nasal delivery of Adeno-Cre that simultaneously activates *KRAS^G12C^*allele and loses *Trp53* or *Lkb1* in K_C_P and K_C_L, respectively and the treatment-naïve tumors uniformly displayed lung ADC pathology (Figure S2A).

We then treated tumor-bearing K_C_L and K_C_P mice with adagrasib (Figure 2A) and found that the tumors were initially responsive to treatment, with maximal tumor regressions ranging 30-75% (Figures 2B-2D). To establish adagrasib-resistant tumors, we performed long-term adagrasib treatment for over 40 weeks (Figure 2D). Notably, we found that the adagrasib-resistant K_C_L tumors displayed squamous pathology including pronounced keratinization. These tumors stained positive for ΔNp63 (p40) (Affandi et al., 2018) and had reduced staining for TTF1 compared to vehicle-treated controls (Figures 2E and 2F). The adagrasib-resistant K_C_P tumors maintained ADC pathology with negative ΔNp63 staining and positive TTF1 staining (Figures 2G and 2H). Previous studies have shown that *Lkb1* deficient tumors had decreased enrichment of NKX motif (Pierce et al., 2021) and increased gene activity score for *Trp63* at baseline when compared to the *Lkb1* wildtype tumors (Figure S2B). These changes in epigenetic landscape may increase the plasticity enabling AST in *Lkb1*-deficient tumors under therapeutic pressure. Resistant K_C_P tumors did not harbor secondary mutations in the *KRAS* allele or in the MAPK pathway (data not shown).

**Figure 2.**
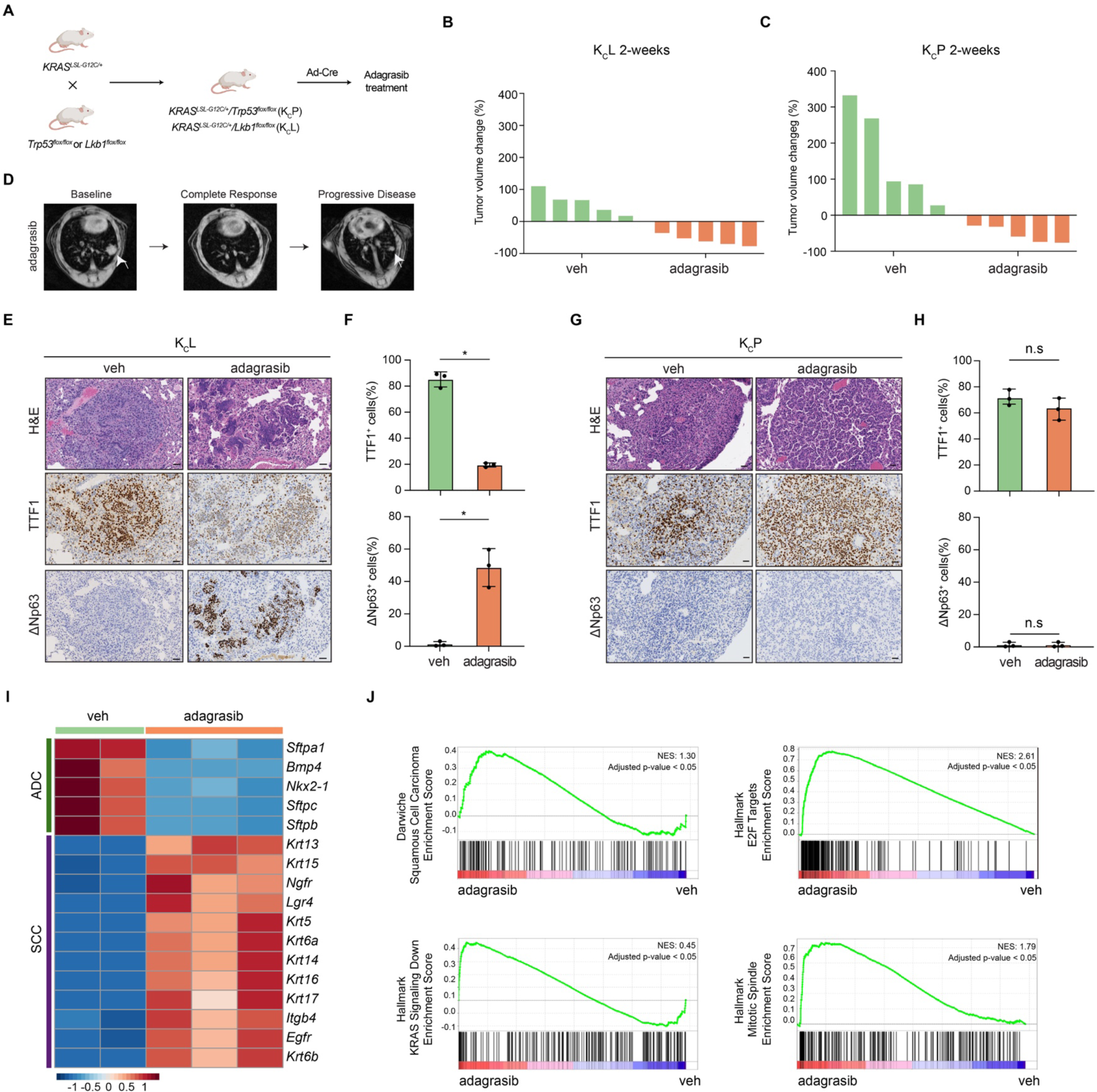
KRAS^G12C^ inhibitor-resistant lung tumors harbor SCC feature. **A.** Schematic illustration of KRAS^G12C^ mouse model study. The K_C_P or K_C_L mice were treated with Ad-Cre and lung tumor burdens were evaluated by magnetic resonance imaging (MRI). Mice were treated with vehicle (veh) (n=3) or KRAS^G12C^ inhibitor adagrasib (100mg/kg) (n=3) for up to 44 weeks until resistance and then subjected to histopathology analyses. **B-C.** Tumor volume changes in K_C_L (**B**) and K_C_P (**C**) mice treated with veh or adagrasib for 2 weeks. **D.** Representative serial MRI photos of K_C_L mice during the acquisition of adagrasib resistance. The white arrows indicate lung tumors. **E.** Representative photos of H&E staining, and TTF1 and ΔNp63 IHC staining in veh-treated and adagrasib-resistant K_C_L tumors. Scale bar, 50μm. **F.** Statistical analyses of the percentage of TTF1 positive cells and ΔNp63 positive cells in veh-treated and adagrasib-resistant K_C_L tumors. * p <0.05. **G.** Representative photos of H&E staining, and TTF1 and ΔNp63 IHC staining of veh-treated and adagrasib-resistant K_C_P tumors. Scale bar, 50μm. **H.** Statistical analyses of the percentage of TTF1 positive cells and ΔNp63 positive cells in veh-treated and adagrasib-resistant K_C_P tumors. n.s = non-significant. **I.** Heatmap and hierarchical clustering of ADC- and SCC-related genes from veh-treated and adagrasib-resistant K_C_L tumors. **J.** GSEA analysis of RNA-seq data for squamous, Hallmark KRAS signaling down and Hallmark proliferation-related signaling pathways.

Consistent with a subset of patients with unknown mechanisms of resistance (Awad et al., 2021), the mechanism of resistance for these K_C_P tumors remains unclear and warrants further studies. Together, these data suggest that AST is a mechanism of acquired resistance to adagrasib that can be modeled preclinically in K_C_L tumors.

To further characterize the K_C_L adagrasib-resistant tumors, we performed transcriptomic analysis of vehicle-treated and adagrasib-resistant K_C_L tumors and found a shift in expression of ADC and SCC lineage-specific genes. Consistent with the tumor histology, vehicle-treated tumors had high expression of ADC-specific genes, such as *Sftpc* and *Nkx2-1*, whereas adagrasib-resistant tumors had high expression of the SCC-specific genes, such as *Krt5* and *Krt14* (Figure 2I). Gene Set Enrichment Analysis (GSEA) of differentially expressed genes (DEGs) between adagrasib-resistant and vehicle-treated tumors revealed significant enrichment of SCC and proliferation-related signatures in adagrasib-resistant K_C_L tumors (Figure 2J). Notably, adagrasib-resistant tumors (collected while on therapy) still displayed down-regulation of KRAS signaling, as indicated by the enrichment of Hallmark ‘KRAS Signaling Down’ signature (Liberzon et al., 2015) (Figures 2J and S2C). Consistently, we found adagrasib-resistant nodules had lower downstream KRAS signaling measured by phosphorylated ERK (pERK) expression relative to vehicle-treated control tumors (Figure S2D). These findings suggest that these adagrasib-resistant tumors were capable of growth *in vivo* despite continued suppression of the RAS signaling pathway by the drug.

Furthermore, to validate AST as a mechanism of resistance to KRAS^G12C^ inhibition, we established an organoid system from K_C_L mice. We harvested tumor nodules from treatment-naïve K_C_L mice for organoid culture according to previously established protocols (Gao et al., 2014a) and continuously treated them with adagrasib (500nM) for 12 weeks (Figure S3A). Consistent with our *in vivo* observations, adagrasib-treated organoids displayed increased ΔNp63 staining and decreased TTF1 staining compared to control organoids (Figures S3B-D). The ΔNp63 positive organoids were more resistant to adagrasib when compared to the control organoids not expressing ΔNp63 (Figure S3E).

Together, the data from *in vivo* and *in vitro* models support that K_C_L tumors invoke plasticity programs and undergo AST to transition to a squamous, KRAS-independent state conferring adagrasib resistance.

### *Lkb1*-deficient ADC revealed two subtypes with differential AST potential and response to KRAS inhibition

We have previously reported that lung tumors from *Kras^LSL-G21D/+^/Lkb1^flox/flox^*(K_D_L) mice show spontaneous AST without therapeutic intervention (Han et al., 2014). To further deconvolve the relationship between *Lkb1*-driven lineage plasticity and therapeutic resistance, we established organoids from K_D_L ADC tumors for long-term culture and investigated the histopathology of the organoid-derived allografts (Figure 3A). *In vitro*, we found more than half of the ADC organoids (7/13) gradually changed morphology from vacuolated to solid spheres, and such change co-occurred with increased expression of ΔNp63 with serial passaging (Figures 3B, 3C, S4A and Table S4). To test whether morphological alterations corresponded to histological transition *in vivo*, we transplanted these morphologically changed organoids into mouse lungs and found that the resulting allograft tumors displayed SCC features with notable intercellular bridging and keratinization, in contrast to parental tumors showing ADC pathology (Figures 3B, 3C, S4A and S5A). These allograft tumors expressed squamous markers including SOX2, ΔNp63, KRT14 and KRT5 similar to the K_D_L SCC and human SCC (hSCC) (Figure S5B). On the other hand, the remainder of the ADC organoids (6/13) retained the vacuolar morphology and the corresponding allograft tumors displayed ADC pathology (Figures 3D, S4B and S5C). Transcriptomic analysis revealed that the organoids with squamous transition expressed higher levels of SCC-specific genes and lower levels of ADC-specific genes (Figure 3E), and were enriched for SCC and proliferation-related signatures (Figure 3F and S6A). Thus, we have developed organoid models to study the relationship between AST and therapeutic intervention, defining a set of mouse K_D_L lung ADC organoids that are highly plastic and capable of undergoing AST (i.e. “plastic” organoid) as well as a comparator set of K_D_L lung ADC organoids that retain ADC features and do not show this plasticity (i.e. “non-plastic” organoids).

**Figure 3.**
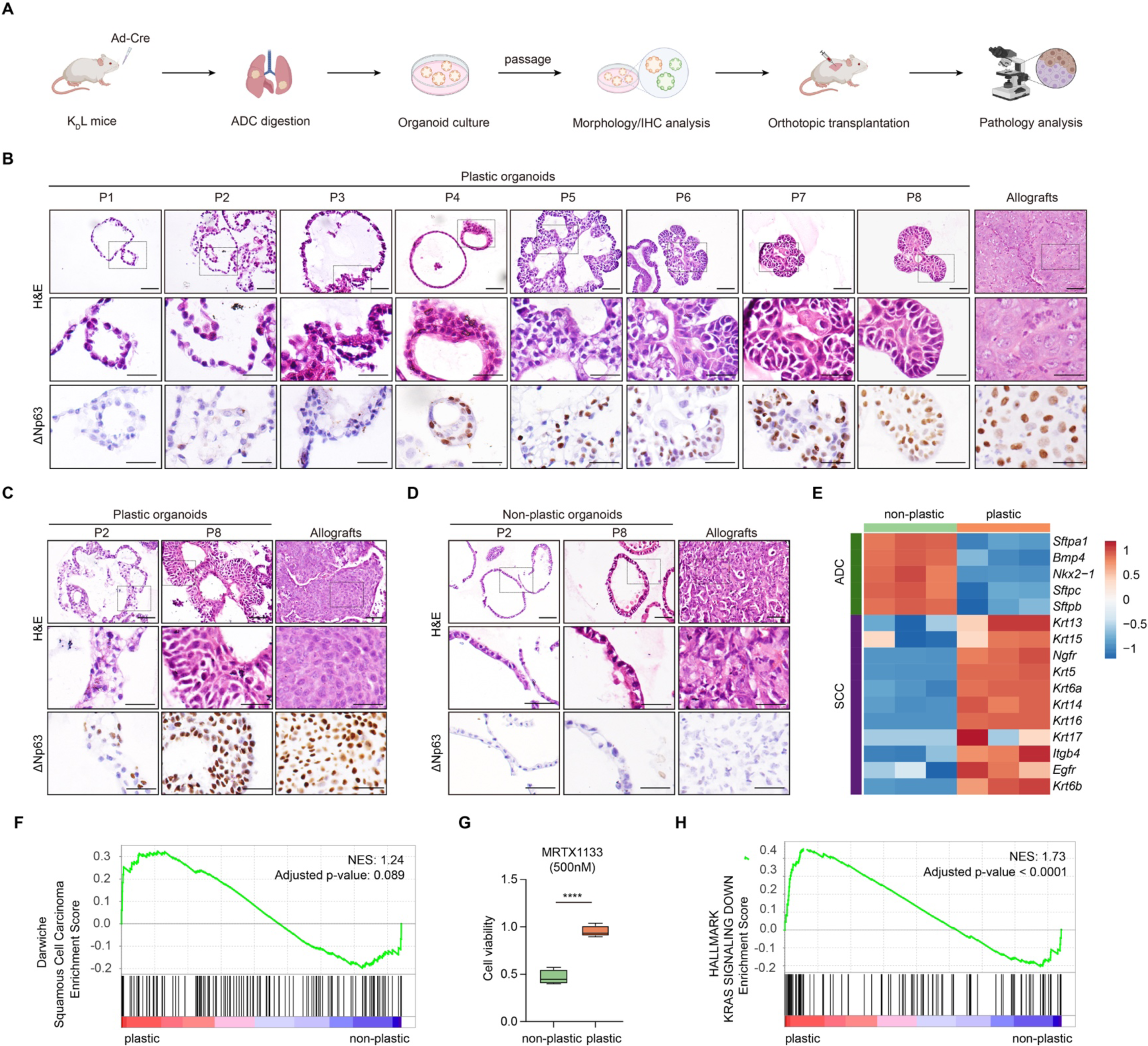
Identification of two different subtypes of K_D_L ADC organoids with differential AST potentials and responses to KRAS inhibition. **A.** Schematic illustration of experiments. The K_D_L lung ADC was collected for the establishment of organoids culture, which was then subjected to orthotopic transplantation and tumor pathological inspection. **B.** Representative photos of H&E staining and ΔNp63 immunostaining for plastic organoids (PO-1) at indicated passages and the corresponding allografts (n=3 mice). Scale bar, 50μm (top panel), 25μm (middle and bottom panels). **C.** Representative photos of H&E staining and ΔNp63 immunostaining for plastic organoids (PO-3) at indicated passages and the corresponding allografts (n=2 mice). Scale bar, 50μm (top panel), 25μm (middle and bottom panels). **D.** Representative photos of H&E staining and ΔNp63 immunostaining for non-plastic organoids (NPO-1) at indicated passages and the corresponding allografts (n=3 mice). Scale bar, 50μm (top panel), 25μm (middle and bottom panels). **E.** Heatmap showing differential expression of ADC- and SCC- related genes in non-plastic and plastic organoids. **F.** GSEA analysis of RNA-seq data for Darwiche Squamous Cell Carcinoma Signature in plastic and non-plastic organoids. **G.** Cell viability of non-plastic and plastic organoids treated with MRTX1133 (500nM) for 3 days. **H.** GSEA analysis of RNA-seq data for Hallmark KRAS signaling down pathway in plastic and non-plastic organoids.

Then, we compared the responses of the plastic and non-plastic organoids to the KRAS^G12D^ inhibitor, MRTX1133 (Figure S6B). Notably, we found the plastic organoids were relatively more resistant to MRTX1133 treatment compared with the non-plastic organoids (Figures 3G and S6B). Immunoblotting revealed pERK was downregulated in MRTX1133 treated organoids and confirmed MRTX1133 was inhibiting KRAS signaling in both plastic and non-plastic organoids (Figures S6C and S6D). These data imply that the plastic organoids may not depend as much on KRAS signaling for proliferation and survival compared with non-plastic organoids, thus promoting the resistance of these cells to KRAS inhibition. Consistently, GSEA revealed the plastic organoids were enriched for ‘Hallmark ‘KRAS Signaling Down’ signature (Liberzon et al., 2015) in the absence of drug treatment (Figure 3H).

Furthermore, we harvested ADC and SCC tumors from the K_D_L GEMM and generated organoids (Figure S7A). We validated K_D_L ADC organoids were positive for TTF1 staining and were negative for ΔNp63 expression while K_D_L SCC organoids were positive for ΔNp63 staining and negative for TTF1 expression (Figure S7B). We then tested the sensitivity of both K_D_L ADC and SCC to KRAS inhibitor. We found that MRTX1133 treatment significantly inhibited the cell viability of K_D_L ADC organoids but had no effect on K_D_L SCC organoids (Figures S7C and S7D). These results support our observation that tumors and organoids that have undergone AST are resistant to KRAS inhibitor treatment.

Collectively, we demonstrate that *Kras*;*Lkb1* mutant ADC harbors a propensity for lineage plasticity and AST is associated with poor response to KRAS inhibitor. This is consistent with the clinical observations that *KRAS*;*LKB1* mutant ADC with high SCC signature at baseline had poor response to KRAS inhibitor treatment.

### Epigenetic regulation of **Δ**Np63 expression drives AST and resistance to KRAS inhibition

To define the underlying mechanisms of AST, we performed integrative analyses of bulk RNA-seq and assay for transposase-accessible chromatin sequencing (ATAC-seq) in plastic organoids undergoing AST during serial passages (Figure 4A). To identify passage-specific genes, we used an entropy-based method (Xie et al., 2013) and found the expression of passage-specific genes were dynamically changed during AST (Figure 4B). We also identified the overlapping passage-specific genes using ATAC-seq data (Figure 4C and Table S5). We found several interesting TFs at specific passages, including *Elf5*, *Atoh8*, *Hmgn3* at early passages, and *H1fx* at intermediate passage. *Elf5* is highly expressed in glandular or secretory epithelial cells (Chakrabarti et al., 2012) and we found that *Elf5*, *Scgb1a1* and *Spdef* were all enriched at early passages, indicating a potential gastric differentiation at initiation (Snyder et al., 2013). Previous studies have shown *Atoh8* is involved in SMAD signal transduction, which might also affect the cancer plasticity (Gabitova-Cornell et al., 2020). *Hmgn3* modulates the structure of chromatin and enhances the transcription from chromatin templates (Reeves, 2015). H1fx is a linker histone that play important roles in DNA damage repair (Hyslop et al., 2005; Valencia and Kadoch, 2019). We speculate that these two genes might play roles in controlling the gene expression changes during AST.

**Figure 4.**
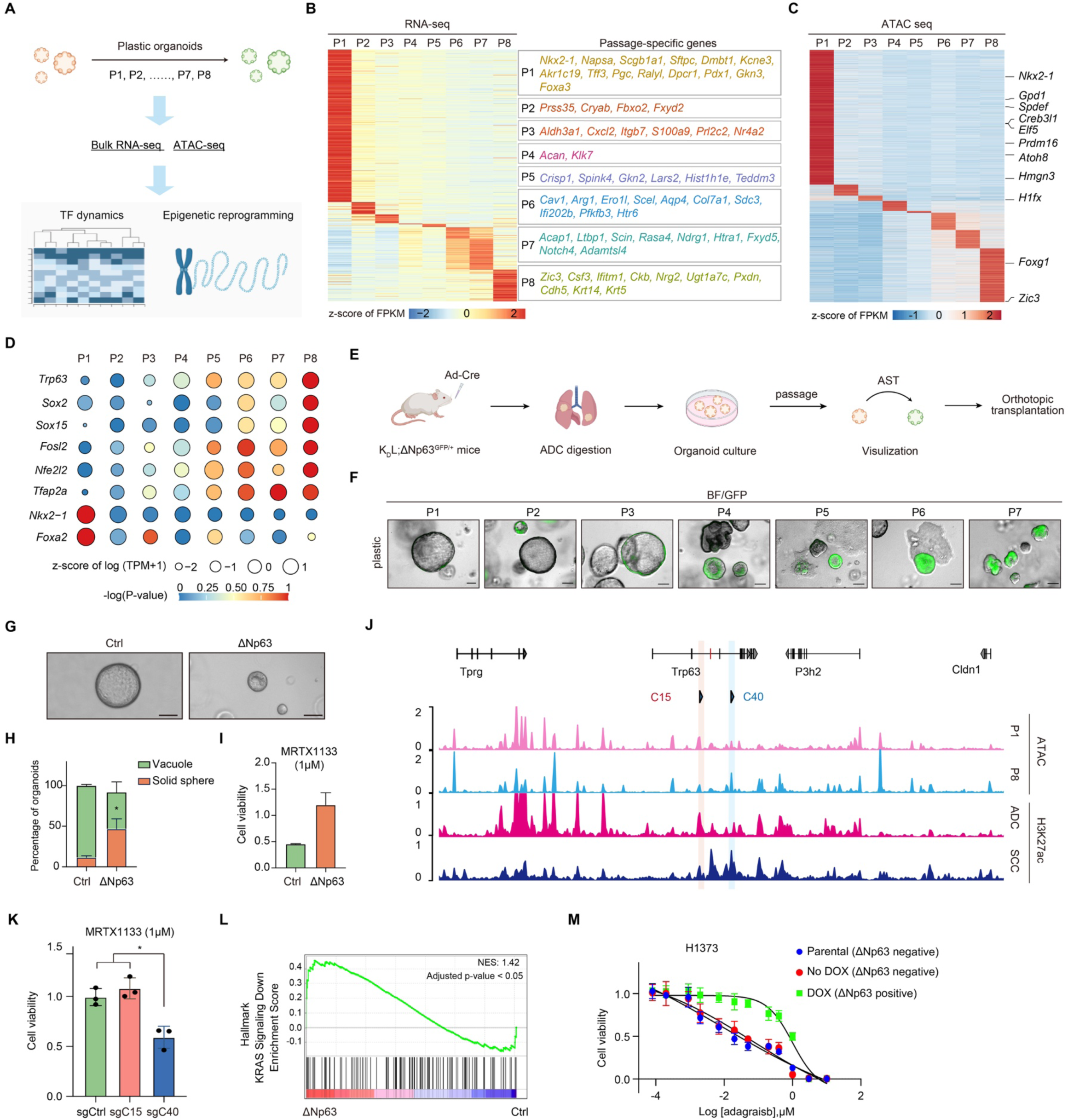
Epigenetic analyses uncover the important role and regulatory mechanism of ΔNp63 in squamous transition and KRAS inhibitor resistance. **A.** Schematic illustration of epigenetic analyses study. **B.** Heatmap visualization of the expression of passage-specific genes from bulk RNA seq. **C.** Heatmap visualization of the ATAC-seq enrichment of passage-specific promoters (TSS±2.5kb). Representative TFs were listed on the right. **D.** Gene expression and enrichment of TF motifs identified from genome-wide ATAC-seq peaks with the passaging. **E.** Schematic illustration of K_D_L;p63^GFP/+^ (K_D_L-p63) mouse model study. **F.** Representative merged photos of bright-field (BF) and fluorescence (GFP) of plastic organoids with the passaging. Scale bar, 50μm. **G-H.** Representative bright-field photos of ctrl and ΔNp63 overexpression organoids (**G**) and statistical analysis of morphology alteration (**H**). **I.** Cell viability of ctrl and ΔNp63 overexpression organoids treated with MRTX1133 (1μM) for 3 days. **J.** ATAC-seq profiles show the chromatin accessibility around *Trp63* loci (C15 enhancer, red box; C40 enhancer, blue box) of plastic organoids at P1 and P8. ChIP-seq profiles (Zhang et al., 2017) show the signals of H3K27ac around *Trp63* loci in K_D_L ADC and SCC. **K.** Cell viability of Ctrl, sgC15 and sgC40 organoids treated with MRTX1133. **L.** GSEA analysis of RNA-seq data for Hallmark KRAS signaling down pathway in Ctrl and ΔNp63 overexpression H1373 cells. **M.** Cell viability of parental, no doxycline treated (ΔNp63 negative) and doxycline treated (ΔNp63 positive) H1373 cells with adagrasib treatments.

The ADC-lineage transcription factors (TFs) including *Nkx2-1* were mainly expressed at early passages concomitant with increased chromatin accessibility at their promoters (Figures 4B and 4C). Conversely, the expression and the chromatin accessibility at promoters of SCC-related genes such as *Krt14* were mainly enriched at late passages (Table S5). Motif analyses using HOMER (Heinz et al., 2010) further revealed that the motif enrichment of ADC lineage-related TFs (*e.g. Foxa2*, *Nkx2-1*) were enriched at earlier passages, while the motifs of SCC lineage-related TFs (including *Trp63* and *Sox2*) demonstrated enrichment at later passage, supporting a crucial role for these TFs in regulating squamous transition (Figure 4D). Moreover, we analyzed the relative changes in expression of lineage-specific TFs in plastic and non-plastic organoids between P4 and P8 (Figure S8A), and found that Δ*Np63* expression increased over passages in plastic organoids, but the expression of *Sox2* was rather stable during passages, potentially suggesting a more prominent role of Δ*Np63* as a driver of AST in these plastic organoids.

To validate the enrichment of Δ*Np63* motif amongst accessible regions and its expression during AST, we crossed the K_D_L mice with Δ*Np63^GFP/+^* mice, in which GFP is knocked-in to the endogenous Δ*Np63* locus in one allele and marks Δ*Np63* expression (Romano et al., 2012). Thus, these K_D_L;p63^GFP/+^ (K_D_L-p63) mice allowed us to generate lung cancer organoids in which continuous monitoring of GFP expression from the Δ*Np63* locus can be performed (Figures 4E and S8B) (Romano et al., 2012). Comparable to K_D_L lung cancer organoids, 50% (3/6) of the K_D_L-p63 ADC organoids switched morphology from vacuolated to solid spheres with serial passaging (Figure 4F and Table S6). Within the plastic K_D_L-p63 organoids the proportion of GFP-positive cells gradually increased with the passaging (Figure S8C). In contrast to plastic organoids, the non-plastic organoids remained vacuolated and were negative for GFP (Figures S8D). Allograft tumors from the plastic organoids showed squamous histology and stained positive for ΔNp63 (Figure S8E) whereas those from the non-plastic organoids retained ADC histology and were negative for ΔNp63 (Figure S8F).

Next, we aimed to test whether ΔNp63 expression is required for maintenance of SCC features and altered response to KRAS inhibition. We overexpressed ΔNp63 in K_D_L organoids at early passages and found a 20∼60% increase of organoids with solid sphere morphology (Figure 4G and 4H). The overexpression of ΔNp63 in non-plastic K_D_L led to increase the expression of keratin related gene *Krt6a* (Figure S9A). To investigate the role of ΔNp63 in modulating response to MRTX1133, we tested for the sensitivity of the organoids and found that ΔNp63 overexpression promoted resistance to KRAS inhibitor MRTX1133 (Figure 4I).

We speculated that ΔNp63 expression was regulated at the epigenetic level in plastic organoids by differential enhancer activity between early- and late- passage plastic organoids that induced the expression of Δ*Np63* during AST as indicated by ATAC and H3K27ac signals (Zhang et al., 2017). Previous work has shown that C15 (359bp) and C40 (334bp) are two conserved enhancer modules at *Trp63* gene body (Antonini et al., 2006). We observed rewiring of the chromatin landscape at *Trp63* locus, with reduced chromatin accessibility at the C15 enhancer, as well as increased accessibility at the C40 enhancer in late-passage plastic organoids compared to early-passage plastic organoids (Figure 4J). Consistently, we found increased H3K27ac signals at the C40 enhancer in K_D_L SCC tumors compared with K_D_L ADC tumors (Zhang et al., 2017). We found that C40 knockout but not C15 knockout induced a loss of solid sphere morphology and decreased ΔNp63 expression in plastic organoids (Figures S9B-E). Notably, we found that the C40-knockout organoids showed an increased sensitivity to MRTX1133 (Figure 4K). In contrast, the control and C15-knockout organoids were resistant to the drug. These results imply that ΔNp63 is important for the maintenance of SCC-like morphology and resistance to KRAS inhibition.

To test whether a squamous state induced by ΔNp63 overexpression is sufficient to induce adagrasib resistance in human lung ADC, we overexpressed ΔNp63 in NCI-H1373, a KRAS^G12C^-mutant human ADC cell line (Figure S9F). Overexpression of ΔNp63 led to the increase in expression of known-squamous related genes (Figure S9G). GSEA analysis revealed ΔNp63 overexpressed cells were enriched for gene-ontology modules such as ‘apical junction’, indicative of a change in cell identity (Figure S9H). Notably, these cells were enriched for ‘KRAS Signaling Down’ suggesting that the activation of ΔNp63 results in inducing a KRAS-independent state (Figure 4L). To test the response to KRAS inhibition, we then treated the ΔNp63-expressing and control cell lines with adagrasib and observed a shift in the drug response-curve with adagrasib treatment upon ΔNp63 induction relative to adagrasib-sensitive control cells (Figure 4M). Together, these data suggest that ΔNp63 overexpression in human lung ADC cells reduces KRAS dependence and confers resistance to KRAS inhibition.

### Single cell RNA-seq analysis reveals squamous transition trajectory and the intermediate state with high plasticity

To investigate cell-fate changes during AST, we performed single-cell RNA-sequencing (scRNA-seq) on plastic organoids at different passages (12,839 cells from P1, P3 and P8) (Figure 5A) and classified these cells into 12 clusters (Figures 5B and S10A). To annotate clusters to a cell state, we examined the enrichment of gene modules in plastic organoids using previously established signatures of lung ADC and SCC (Inamura et al., 2005; Marjanovic et al., 2020) (Figure 5C). We found cells in clusters 1-3 were enriched for alveolar cell type 2 (AT2) gene module with high expression of *Sftpb*, suggesting that these cells are in ADC state given that lung ADC predominantly arises from a subset of AT2 cells (Desai et al., 2014; Sutherland et al., 2014). Additionally, we found the SCC signature was enriched in clusters 10-12, which was mainly from the later passage (P8) and expressed canonical SCC marker genes such as *Krt5* and *Krt14* (Figures 5C and S10B). We found that clusters 4-6 expressed gastrointestinal epithelium (GI)-like signature, clusters 6-7 expressed EMT signature and clusters 8-9 expressed embryonic liver-like and gastric-like signatures which were not enriched for AT2 or SCC gene modules. These results indicated that clusters 4-9 expressed features of different cell types, suggesting the potential for high cellular plasticity. Indeed, the intermediate clusters 4-7 were enriched for the high-plasticity cell state (HPCS) signature which was previously shown to be associated with high potential for cell state transition (Marjanovic et al., 2020) (Figure 5D). To further identify the cellular identity of the HPCS, we computed scores of several reported lineage-related gene modules, including alveolar, basal and glandular cells (Barkley, 2022) (Figures 5E and S10B). We found that glandular modules were uniformly expressed across the three passages (P1, P3, P8). In intermediate clusters 4-7 with high HPCS signatures, the cells co-expressed alveolar and basal-related signatures (Figure 5E).

**Figure 5.**
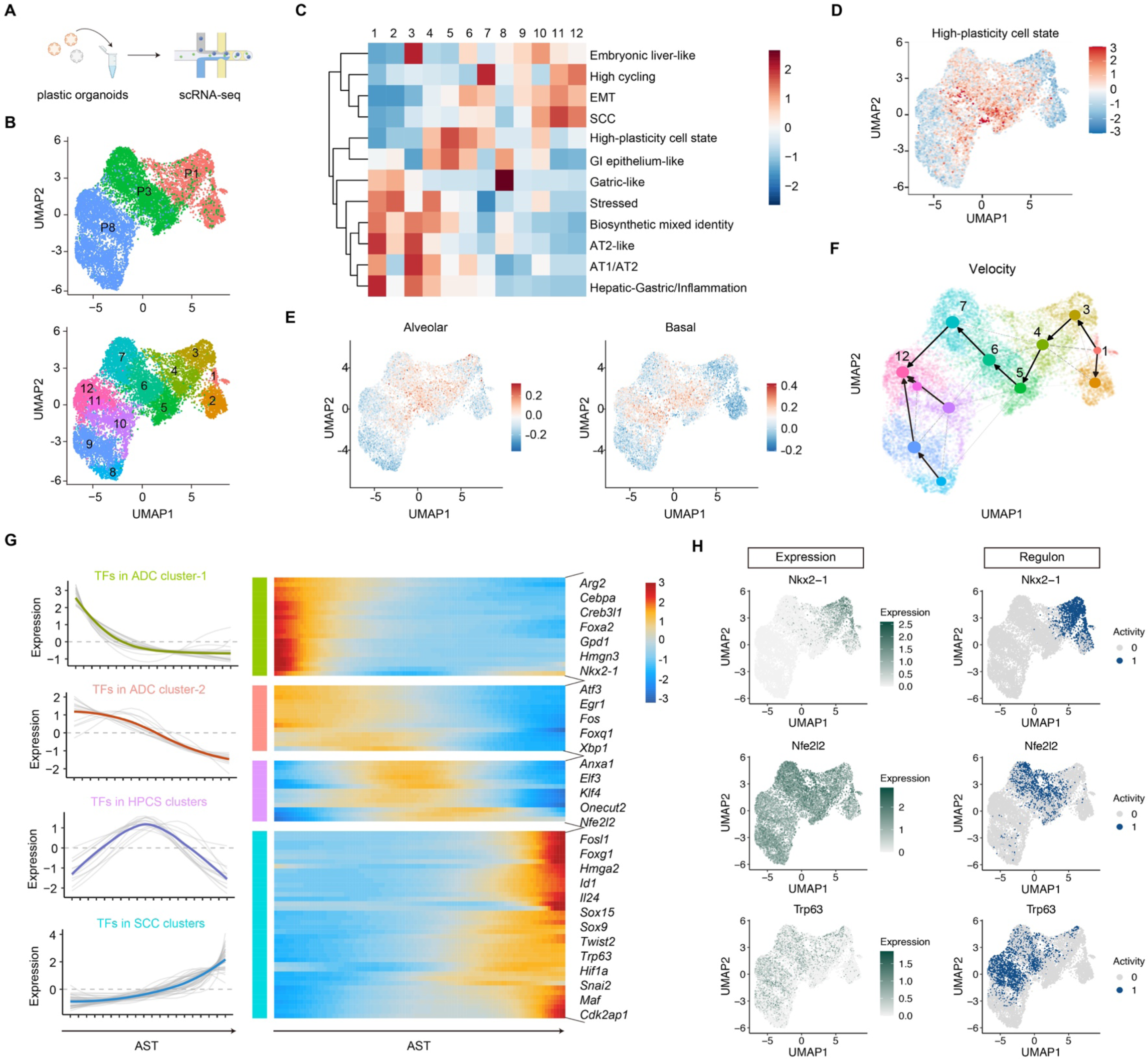
Identification of an intermediate state of K_D_L organoids during AST featured with a high-plasticity cell state signature. **A.** Schematic illustration of single-cell transcriptome analyses of organoids. **B.** Uniform Manifold Approximation and Projection (UMAP) visualization of organoids at single-cell level with the passaging (P1, P3, P8). **C.** Heatmap showing signature scores in different clusters annotated based on Marjanovic et al., 2020. **D.** UMAP visualization of AUC scores of high-plasticity cell state (HPCS) signatures. **E.** UMAP visualization of alveolar and basal cell modules scores as a function of average expression of module genes relative to background from <u>Barkley, 2022</u>. **F.** Partition-based graph abstraction (PAGA) maps for indicated clusters. Weighted edges correspond to the connectivity between two clusters, and arrows to velocity-inferred directionality. **G.** Gene trends for TFs across the AST branch probability. Gene trends were grouped into four clusters. Solid and dashed lines represent the mean and standard deviation of the gene trends for each cluster. **H.** UMAP visualization of expression level (left panel) and binary target gene set (regulon) activity (right panel) of *Nkx2-1*, *Nfe2l2*, Δ*Np63*.

To analyze the temporal sequence of states of individual cells during AST, we performed RNA velocity analysis and Partition-Based Graphical Abstraction (PAGA) (Bergen et al., 2020; La Manno et al., 2018; Wolf et al., 2019). The trajectory analyzed by this method showed clusters 1 and 3 exhibiting positive velocity toward the intermediate clusters 4-7, and eventually entering into cluster 12 (Figures 5F). Integrated analysis of the velocity and the gene modules revealed that the AT2 cells transitioned into a cell type with co-expression of alveolar and basal cell signatures, followed by transition to squamous cells. We also observed that cells in cluster 2, mostly comprised of cells from P1, and cluster 8, mostly comprised of cells from P8, both expressed gastric-like and glandular signatures, indicating some cell states can recur across the passages.

Overall, we show that ADC cells from plastic organoids go through an intermediate state with increased HPCS signatures and an ability to transition to SCC. Interestingly, we also observed that HPCS signature was enriched in one EGFR-mutant lung cancer patient with demonstrated AST upon relapse from osimertinib treatment (Maynard et al., 2020) (Figures S10C), indicating that our model might recapitulate plasticity features of drug-induced AST patients.

Next, we aimed to identify TFs that might be responsible for the lineage plasticity. First, we defined differentially enriched TFs ordered by pseudotime, and then TFs were grouped into four major trends. These groups included TFs expressed in ADC clusters (2 groups), in HPCS clusters and in SCC clusters (Figure 5G). The expression of TFs in ADC clusters quickly (cluster-1) or gradually (cluster-2) decreased over the passages, the TFs in HPCS clusters were mainly enriched at intermediate state and the expression of TFs in SCC clusters was gradually increased during AST. To investigate the activity of these TFs in driving squamous transition in plastic organoids, we performed gene regulatory network inference using the SCENIC algorithm (Aibar et al., 2017). SCENIC identifies a TF-regulon that is a group of co-expressed genes inferred to be regulated by a specific TF and computes regulon activity based on the level of expression of its member genes compared to all other genes. We found the regulon activity of *Nkx2-1*, whose expression is mainly enriched in ADC clusters, was low in clusters 4-7, suggesting a loss of ADC features (Figure 5H). We observed the expression and regulon activity of *Nfe2l2* was high in HPCS clusters 5-7, indicating a potential function of *Nfe2l2* regulating oxidative stress in AST plasticity. Notably, the regulon activity of Δ*Np63* was high in clusters 6 and 7 (Figure 5H), indicating that there was a group of genes activated by Δ*Np63* in the HPCS. The Δ*Np63* regulon activity was maintained in SCC clusters 10-12 and includes several squamous-related genes, include *Krt5* (Table S7). These results suggest that Δ*Np63* and associated gene expression changes during HPCS drive the AST and maintain the SCC features.

The GO biological pathway analysis of the DEGs along the pseudotime identified several enriched pathways, including metabolic, ECM remodeling, chemokine, and proliferation related pathways (Figure S10D). The metabolic processes, including fatty acid metabolism and oxidative phosphorylation, were gradually decreased during AST. The signaling enrichment of cell junction organization and keratinocyte differentiation was increased in SCC clusters. We noticed that leukocyte chemotaxis was increased during AST, suggesting the intrinsic factors might regulate the microenvironment to affect AST. At the end of the trajectory, genes of cell cycle and division related pathways were activated, indicative of highly proliferative property of SCC.

### *KRT6A* expression correlates with adagrasib resistance in *KRAS*;*LKB1* mutant NSCLC

To identify markers of squamous transition and resistance to adagrasib, we defined commonly enriched genes in HPCS clusters 6 and 7, the Δ*Np63* regulon and adagrasib-resistant tumors. We found six overlapping genes marking an adagrasib-resistant, ‘AST plasticity’ signature, including *Krt6a*, *Aqp3*, *Wnt4* and *Sfn* (Figure 6A).

**Figure 6:**
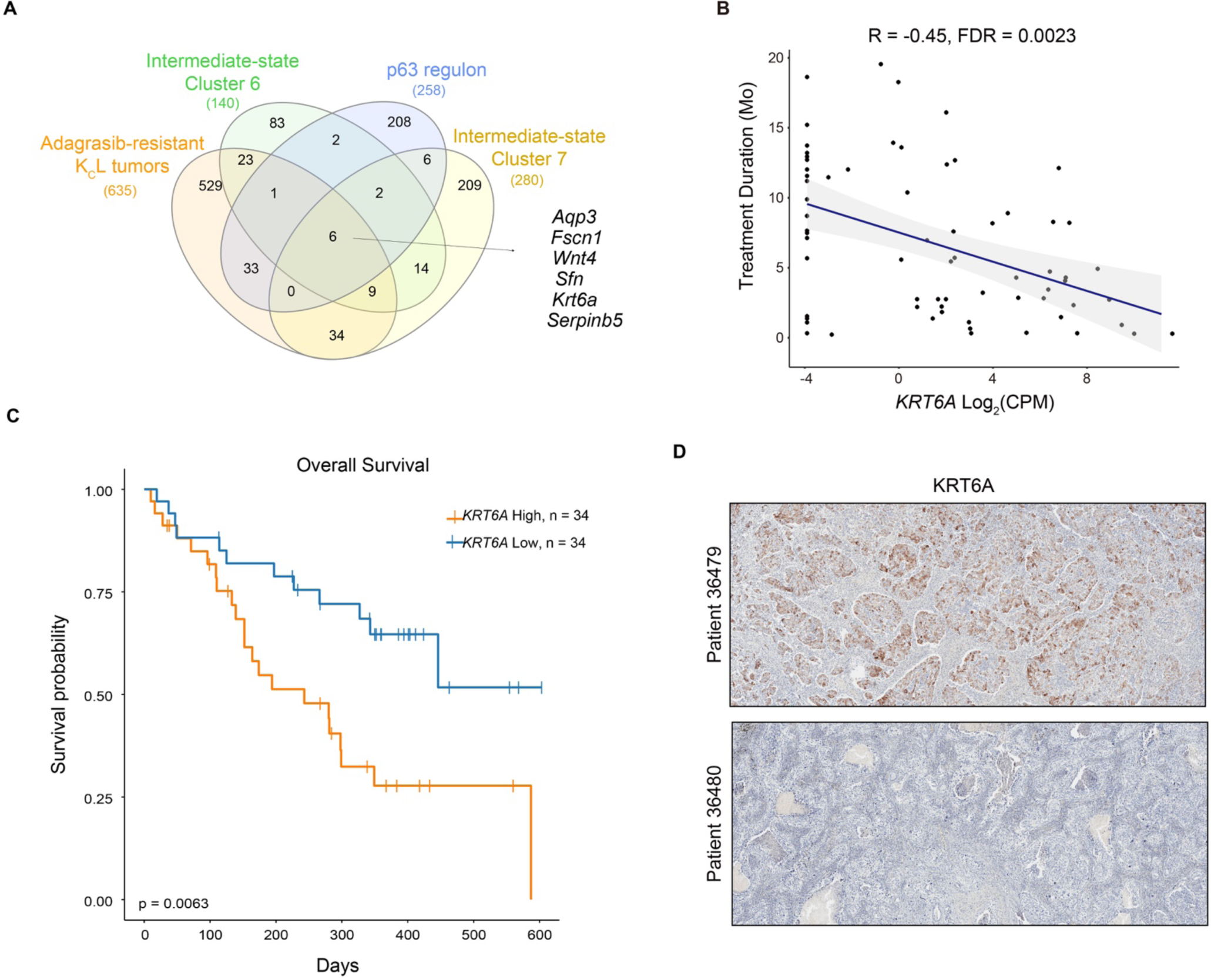
KRT6A gene expression in pre-treatment biopsies associate with adverse outcomes to adagrasib monotherapy in lung cancer. **A.** Overlap analysis of enriched genes in adagrasib-resistant K_C_L tumors, Δ*Np63* regulon and the intermediate state (including clusters 6 and 7). **B.** Scatter plots showing correlation between *KRT6A* expression and time on treatment. **C.** Kaplan-Meier curves showing probability of survival for patients stratified by *KRT6A* expression above the median (high) and below the median expression (low). **D.** Representative immunostaining photos for tumors positive and negative for KRT6A in an independent cohort of surgically resected lung adenocarcinomas without previous treatments.

To determine how the AST plasticity signature links to response to adagrasib treatment, we correlated the expression level of *KRT6A* with the treatment duration in patients from KRYSTAL-1 trial. We found that expression of *KRT6A* showed a statistically significant negative correlation with treatment duration upon correction for multiple hypothesis testing (Figure 6B and Table S3). When we stratified patients above or below the median *KRT6A* expression (i.e., high- and low- *KRT6A* expression), we observed that patients with high expression of *KRT6A* had a significantly shorter OS and PFS than those with low expression (Figures 6C and S11A). Patients with high *KRT6A* expression also had statistically a significant higher frequency of *KEAP1* and *STK11*/*LKB1* mutations. *KEAP1* mutations have been shown to correlate with a lower response rate to adagrasib therapy (Figures S11B and S11C) (Jänne et al., 2022; Skoulidis et al., 2021). When tumors were *KEAP1* wildtype, *KRT6A* expression still demonstrated a negative association with adagrasib treatment duration in both *STK11*/*LKB1* wildtype and mutant genotypes (Figures S11D and S11E). These data suggest that *KRT6A* expression correlates with worse outcomes on adagrasib monotherapy in NSCLC patients.

We examined the expression of KRT6A in mouse tumors and found that KRT6A was expressed in a subset of K_D_L ADC cells, but rarely detected in K_D_P ADC. KRT6A was also highly expressed in spontaneously transitioned K_D_L SCC (Figure S12). The results were consistent with our finding that *Krt6a* expression preceded the appearance of SCC signature, and hinted that some *Kras*;*Lkb1* ADC with KRT6A expression might harbor potential AST plasticity. Furthermore, in an independent cohort of NSCLC patients we identified 5/36 patients expressing KRT6A by immunohistochemistry (Figure 6D), suggesting this might demark a subset of patients that harbor inherent lineage plasticity that may become resistant to RAS-targeted therapy through AST.

## Discussion

In this study, we examined gene expression features and clinical outcomes from a cohort of NSCLC patients treated with adagrasib monotherapy from the KRYSTAL-1 trial, and we demonstrated that the enrichment of SCC signature in the pre-treatment biopsies was correlated with adverse clinical outcomes, specifically in patients with co-occurring *KRAS*^G12C^ and *STK11*/*LKB1* mutations. We further showed that long-term adagrasib treatments of K_c_L (*KRAS^LSL-G12C/+^/Lkb1^flox/flox^*) GEMMs and organoids led to acquired resistance, with tumors from the K_c_L model demonstrating striking enrichment of SCC feature at both histologic and molecular levels. Squamous transition has been proposed as an emerging mechanism for molecular targeted therapy resistance, e.g., EGFR TKI resistance (Chen et al., 2019; Cooper et al., 2022; Passaro et al., 2021; Schoenfeld et al., 2020). However, it remains largely unknown about its potential contribution to KRAS inhibitor resistance. We recently reported two KRAS^G12C^ lung ADC patients showing squamous pathology after relapse (Awad et al., 2021). This observation triggered us to investigate whether squamous transition is a collateral event or directly drives drug resistance. Our current study combining mouse and human studies strongly support the causality between AST and KRAS inhibitor resistance; squamous transition potentially endows lung adenocarcinoma with inherent plasticity and make them less dependent on the KRAS signaling and thus become resistant to KRAS inhibitors. Moreover, through integrative analyses of the evolutionary mechanisms of squamous transition using mouse models and clinical data, we identified an intermediate state during transition featured with high-plasticity cell state signature marked by *Krt6a* upregulation. Furthermore, *KRT6A* expression is significantly correlated with poor clinical responses to adagrasib in KRAS^G12C^-mutant lung cancer. These findings might bring immediate benefit for clinical prediction of KRAS inhibitor efficacy.

Our clinical data implicate that AST is significantly associated with adagrasib resistance in the context of *STK11*/*LKB1* mutations. A recent study reported squamous transition in relapsed ADC patients mainly receiving EGFR TKI and its relationship with *STK11*/*LKB1* mutations (Quintanal-Villalonga et al., 2021). Even with its near mutual exclusivity with EGFR mutations (Sanchez-Cespedes et al., 2002; The Cancer Genome Atlas Research Network, 2014), *STK11*/*LKB1* mutation was found in one of three transitioned samples. Additionally, in our own analyses, one of the two relapsed patients with squamous transition harbored a *STK11*/*LKB1* mutation. These clinical observations highlight a potentially important role of LKB1 in squamous transition and drug resistance. Here, we found long-term treatment of adagrasib promotes squamous transition specifically in KcL model but not in K_c_P model, indicative of differential drug resistance mechanisms in *LKB1* or *Trp53* mutation contexts (Salmón et al., 2023). It is worth noting that both K_C_P and K_C_L GEMMs uniformly displayed lung ADC pathology even at humane end-point. In contrast, the K_D_L model exhibited spontaneous AST (Han et al., 2014). This difference may be somehow related to model-specific discrepancy or differential downstream signaling associated with Kras^G12D^ and Kras^G12C^ (Li et al., 2018; Zafra et al., 2020). Nonetheless, we found that the molecular pathways involved in spontaneous AST or drug-induced AST are comparable. For example, similar changes in gene signatures are observed during AST, e.g., the increased proliferation-related signatures and decreased KRAS signature besides the increased SCC signature. Moreover, the upregulation of *Krt6a* expression was also observed in both models. Aligned with our GEMM findings, we also observed an enrichment of high-plasticity cell state in one AST patient relapsed from osimertinib treatment.

We speculate that *STK11*/*LKB1* mutation may induce epigenetic plasticity that enables the AST process. Indeed, in our analysis of the KRYSTAL-1 cohort of human lung ADC patients treated with adagrasib, a subset of *STK11*/*LKB1* mutant tumors were enriched for the SCC signature. Overall, these data suggest that *STK11*/*LKB1* mutant patients harbor inherent plasticity at baseline, with representation of squamous phenotype and its associated SCC signature enrichment predicting a shorter clinical benefit to adagrasib therapy. The *KRAS*-mutant, *STK11/LKB1* mutant subset (i.e. ‘KL’ subtype) of lung adenocarcinoma is a particularly challenging clinical subtype, with poorer responses to standard treatments (Hong et al., 2023; Koyama et al., 2016; Skoulidis et al., 2018). While *STK11/LKB1* mutation itself does not independently stratify responses to KRAS^G12C^ inhibitors in lung cancer patients treated on early clinical trials (Dy et al., 2023; Jänne et al., 2022; Negrao et al., 2023; Skoulidis et al., 2021), the data presented here suggest that the KL subtype can be further subdivided by enrichment of a SCC phenotype and that the KL/SCC-high subtype responds poorly to adagrasib monotherapy. Development of novel combination therapy strategies targeting AST in the KL/SCC-high subtype will be needed to maximize the impact of KRAS inhibition in these patients.

Integrative transcriptomic and epigenomic analyses identified Δ*Np63* as a driver of AST. Modulating the expression of Δ*Np63* altered the response to KRAS inhibition, with overexpression of Δ*Np63* leading to resistance to KRAS inhibition and enhancer knockout at the *Trp63* locus restoring sensitivity to KRAS inhibition. We found that ΔNp63 overexpression upregulated keratin-related genes and squamous-related signatures and modulated cell identity. Moreover, ΔNp63 inhibits the enrichment of ‘KRAS Signaling Down’ signature to mediate poor response to KRAS inhibitors. Using scRNA-seq of plastic organoid models, we further identified the emergence of an intermediate state that was marked by the expression of a plasticity signature and ΔNp63-regulated genes, such as *Krt6a*. We further demonstrated that high *KRT6A* expression in the pre-treatment biopsies was correlated with adverse clinical outcomes. Collectively, we provide preclinical mechanistic data and clinical data to support a prominent role of AST in mediating KRAS inhibitor resistance. Furthermore, our results indicate that *KRT6A* is a general biomarker that may be applied clinically to define a plastic cancer cell state that is predisposed to undergo AST and may predict an adverse outcome with KRAS inhibitor treatment in *KRAS^G12C^*;*LKB1-mutated* context.

We have previously uncovered the epigenetic and metabolic molecular mechanisms underlying the AST process in K_D_L model (Fang et al., 2023; Gao et al., 2014b; Han et al., 2014; Li et al., 2015; Tang et al., 2023). We found that the dynamic dysregulation of the counteracting lineage-specific TFs including ADC related TFs NKX2-1, FOXA2, and SCC related TFs ΔNp63 and SOX2, finely tuned the AST process (Tang et al., 2023). Additionally, loss of the histone modifier polycomb repressive complex 2 (PRC2) activity or EZH2 inhibition promotes epigenetic rewiring and potentiates the AST process (Zhang et al., 2017). At the metabolic level, we observed an excessive accumulation of ROS promoted AST potentially through the dysregulation of AMP-activated protein kinase (AMPK)-mediated fatty acid oxidation (FAO) in lung ADC (Li et al., 2015). Here, we also found oxidative phosphorylation and fatty acid signatures were decreased during squamous transition. Importantly, we identified a new intermediate state enriched for a highly plastic cell state signature with the enrichment of *Nfe2l2* regulon which might regulate ROS mediated AST plasticity and drug resistance. Loss of KEAP1 or activation of NFE2L2 was postulated as a biomarker of resistance to therapy in lung cancer in multiple prior studies (Frank et al., 2018; Hellyer et al., 2019; Homma et al., 2009; Krall et al., 2017). In the context of KRAS inhibitor in lung cancer, *KEAP1* mutated patients had lower response rate (28.6%) when compared to *KEAP1* wildtype patients (51.7%) (Jänne et al., 2022). The mechanism of resistance mediated by *KEAP1* mutation is not clearly delineated. Enhanced handling of ROS upon MAPK pathway inhibition and altered expression of metabolic genes allowing cells to bypass MAPK signaling in *KEAP1* mutated cells were suggested as potential mechanisms (Krall et al., 2017). Our data show that *KEAP1* mutation is correlated with higher expression of *KRT6A* and shorter duration on adagrasib therapy in *KRAS^G12C^* mutated lung cancer. Given the early stage of clinical trials of adagrasib in cancer patients, the availability of patient biopsies with transcriptomic data is relatively limited, and the sample size analyzed in this study is relatively small. The precise relationship of *KRT6A* and squamous-associated gene expression with *KEAP1* mutation status will need to be further dissected in larger cohorts, and also including other KRAS inhibitors such as sotorasib in future studies.

In summary, we provide evidence that AST in lung cancer is strongly associated with KRAS inhibitor resistance in preclinical models and demonstrate that markers of AST plasticity and the SCC phenotype in pre-treatment biopsies may predict poorer responses and reduced clinical benefit from KRAS^G12C^ inhibitors in *KRAS*;*LKB1* mutant lung cancer.

## Methods

### Mouse studies

The *Lkb1^flox/flox^* or *Trp53^flox/flox^* mice were crossed with *Kras^LSL-G12D/+^* mice or *KRAS^LSL-G12C/+^*mice (Li et al., 2018) respectively to generate three different mouse models including *Kras^LSL-G12D/+^*/*Lkb1^flox/flox^* (K_D_L), *KRAS^LSL-G12C/+^/Trp53^flox/flox^* (K_C_P) and *KRAS^LSL-G12C/+^/Lkb1^flox/flox^*(K_C_L). Δ*Np63^GFP/GFP^*mice were kindly provided by Dr. Satrajit Sinha. Δ*Np63^GFP/GFP^* mice were crossed with K_D_L mice to generate K_D_L;p63^GFP/+^ (K_D_L-p63) mice. All mouse work was reviewed and approved by the Institutional Animal Care and Use Committee at NYU School of Medicine and the Institutional Animal Care and Use Committee of Shanghai Institute of Biochemistry and Cell Biology, Chinese Academy of Sciences. All mice were kept in specific pathogen-free environment, and age- and sex-matched animals were randomly grouped for this study. Mice were treated with Ad-Cre virus (2×10^6^ PFU) via nasal inhalation at 6-8 weeks old. Lung tumors were dissected from K_D_L mice, K_D_L-p63 mice and K_C_L mice after 6 weeks to 12 weeks post Ad-Cre infection and cut into small pieces for histological inspection and organoid culture.

The K_C_P and K_C_L mice post 6 weeks of Ad-Cre treatment were given with 100mg/kg adagrasib via oral gavage daily and subjected to MRI using BioSpec USR70/30 horizontal bore system (Bruker) (Tang et al., 2022) every 4 weeks until the acquisition of drug resistance. All mice were then sacrificed for detailed pathological analysis.

### Organoid culture and manipulation

Mouse lung tumors from the K_D_L, K_C_L and K_D_L-p63 mice were dissected and incubated with digestion buffer (advanced DMEM/F12 containing 5 mg/mL Collagenase Type II, 1x P/S, 1mg/ml primocin and 10mM Y-27632 for 45 min at 37°C. Cell suspensions were resuspended in organoid culture medium and mixed with Basement Membrane Extract (BME) (Corning) (1:1), and plated in 24-well plate (5000∼10000 cells/per well). The recipe for the organoid culture medium: advanced DMEM/F12 supplemented with 1x P/S (Invitrogen,15140-122), 1x HEPES (Invitrogen, 15630-056), 1x primocin (Invitrogen, ant-pm-1), 1x B27 supplement (Invitrogen, 17504-044), 1.56mM N-Acetylcysteine (Sigma-Aldrich, A9165-5G), 500nM A-83-01 (Tocris, 2939), 10 ng/ml epidermal growth factor (Corning no. 355056), 10 µM Y-27632 (STEMCELL Technologies, no. 07171), 1x Glutamax (Gibco no. 35050), R-spondin and Noggin (Gao et al., 2014a). The cell suspensions (5x10^5) digested from organoids were orthotopically injected into mouse lung. At every passage, photos were taken before digestion.

Regarding to the KRAS inhibitor treatments *in vitro*, approximately 2,000 cells from organoids were mixed with 5μl Matrigel and seeded in 96-well plates. Cell viability was assayed using CellTiter-Glo 3D (Promega) according to the manufacturer’s instructions following 3 or 5 days of drug incubation, and results were normalized to vehicle controls.

Regarding the C15, C40 knockout and ΔNp63 overexpression experiments, the sgRNAs were designed using optimized CRISPR Design (crispr.mit.edu/). Cloning methods were optimized following established protocols (Ran et al., 2013; Wu et al., 2018). The sequences of all sgRNAs primer pairs are shown in Table S8. Lentivirus was generated by transfection of HEK-293T cells with lentiCRISPRv2 (sgC15, sgC40 or sgCtrl) and the packaging plasmids psPAX2 (Addgene #12260) and pMD2.G (Addgene #12259) using PEI (Polyethylenimine). Once organoids were dissociated, cells were pelleted and resuspended in 250 µL lentiviral solution then incubated at 37°C for 12 hours. Puromycin was added to the media to select the organoids with infection. We pick up single organoids infected with sgC15 and sgC40 or pool organoids infected with sgΔNp63 to do further analysis.

### Hematoxylin and Eosin (H&E), Immunohistochemistry (IHC) and Immunofluorescence (IF) Staining

H&E staining and IHC staining were performed as previously described (Gao et al., 2014b; Han et al., 2014). In brief, mouse lung lobes or organoids were fixed with formalin or 4% paraformaldehyde (PFA) respectively overnight, and dehydrated in ethanol, embedded in paraffin, sectioned (5μm) followed by staining with hematoxylin and eosin. For IHC staining, slides were de-paraffinized in xylene and ethanol, and rehydrated in water. Slides were quenched in hydrogen peroxide (3%) to block endogenous peroxidase activity. Antigen retrieval was performed by heating slides in a microwave for 20 minutes in sodium citrate buffer (pH 6.0) or 1 mM EDTA buffer. The slides were incubated with primary antibodies at 4°C overnight and then analyzed using the SPlink Detection Kits (Biotin-Streptavidin HRP Detection Systems) following the manufacturer’s instructions.

For immunofluorescence (IF) staining, tissues or organoids were fixed in 4% PFA for overnight or 15 minutes respectively, then dehydrated in 30% sucrose solution overnight, embedded in optimal cutting temperature compound (OCT), sectioned (5μm) followed by IF staining. Slides were incubated with primary antibodies in 1% BSA+0.25% Triton X-100 in PBS at 4°C overnight, incubated with second antibodies for 1 hour and DAPI for 10 minutes at room temperature, and then covered with coverslip for imaging. The K_D_L organoids and tissues images were acquired using a Leica TCS SP8 WLL confocal microscope. The K_C_L organoids images were acquired using Zeiss 880 Laser Scanning Confocal Microscope and were processed by FIJI (NIH).

Antibodies used in this study include anti-SP-C (Millipore, AB3786, 1:500), anti-KRT5 (Bioworld, BS1208, 1:250), anti-ΔNp63 (Abcam, ab124762, 1:2000), anti-p40 (Maxim, RMA-1006, 1:250; Abcam, ab203826), anti-SOX2 (Abcam, ab92494, 1:500), anti-KRT14 (Abcam, ab51054, 1:500), anti-TTF1 (Abcam, ab133638, 1:500), anti-KRT6A (Sangon Biotech, D220238, 1:2000).

### Bulk RNA sequencing and bioinformatic analysis

Total RNA were extracted using RNeasy Plus Mini Kit (QIAGEN, cat# 74136). The RNA-seq libraries were constructed using the VAHTS™ mRNA-seq v2 Library Prep Kit for Illumina (NR601, Vazyme). In K_D_L model, the plastic organoids and non-plastic organoids at early passages were collected for bulk RNA seq. In the K_C_L model, tumor nodules from adagrasib-resistant mice (after 40 weeks of treatment) and tumor nodules from vehicle group (at humane end point 12 weeks post-induction) were used for bulk RNA-seq. Sequencing adapter and low-quality bases were trimmed from paired-end reads using Cutadapt v1.18 (Martin, 2011) with parameters “-O 3 -n 5 -m 50 -q 25 --pair-filter=any”. Trimmed reads were aligned to mm10 reference genome using hisat2 v2.1.0 (Kim et al., 2015) with default parameters. Alignments were filtered for unmapped, mapped to multiple positions (MAPQ < 10) and not properly paired reads with SAMtools v1.9 (Li et al., 2009). Alignments were sorted and converted to BAM files with index.

Bigwig files of signal tracks were generated using deepTools bamCoverage v3.2.1(Ramírez et al., 2016) with parameters “*--binSize 10 --normalizeUsing CPM*” to scale the bam files to counts per million mapped reads (CPM) in each 10bp bin across the genome. Replicates were averaged to represent the cell state for visualization. The gene-level expression from RNA-Seq was quantified by HTSeq v0.11.0 (Anders et al., 2015) with parameters “*-f bam -r name -s no -a 10 -i gene_id*” and GRCm38.86 gene annotation. Differentially expressed genes (DEGs) between two passages (P1 and P8) were called by DESeq2 v1.22.1 (Love et al., 2014) with protein coding genes considered significant if the absolute value of log2 fold-change (LFC) was above 1 and the FDR was less than 0.05. And PCA analysis was generated with plotPCA function in DESeq2 package. To report the gene expression level, FPKM (Fragments Per Kilobase per Million) and Transcripts Per Kilobase Million (TPM) (Mortazavi et al., 2008) were used to normalize expression values. Non-overlapping exon length of each gene was calculated from genomic annotation gff files for length normalization. TPMs of the same passage were averaged and log transformed to represent the cell state. GSEA (v3.0) from Broad Institute Platform was perform to pathway enrichment (including established “Darwiche Squamous Signature”) and proliferation signature (“Hallmark E2F Targets”) and p-value < 0.05 was set as statistical significance.

### ATAC sequencing and bioinformatics analysis

ATAC-seq libraries of the time-series plastic organoids were prepared from TruePrep DNA Library Prep Kit V2 for Illumina (Vazyme, TD503). Sequencing adapters and low-quality bases were trimmed from paired-end reads using Cutadapt with parameters “*-O 5 -m 36 -q 25 --pair-filter=any*”. Trimmed reads were mapped to the mm10 reference genome using Bowtie2 v2.3.4.2 (Langmead and Salzberg, 2012) with parameters “*-X 2000 --no-mixed --no-discordant*”. Alignments were filtered for unmapped, mapped to multiple positions or not properly paired fragments. PCR duplicates were removed using Piccard MarkDuplicates v2.18.5 and not used in subsequent analysis.

Bigwig files of signal tracks were generated using deepTools bamCoverage with parameters “--binSize 10 --normalizeUsing CPM *--ignoreForNormalization chrM --extendReads*” for visualization. Peak calling for accessible regions was performed using MACS2 callpeak v2.1.2 (Zhang et al., 2008) using the options “*-g mm -f BAMPE --keep-dup all*” to pile up with fragments defined by paired-end reads. Motif enrichment analysis of regions sets were performed using HOMER v4.9.1 (Heinz et al., 2010) with default parameters and additional options of “-mask” to mask repeats, and motifs with p-value below 1e-10 were considered as significant.

An entropy-based method was used to identify passage-specific genes as previously described (Xie et al., 2013) using averaged FPKM in the same passage. Genes with entropy value less than 2.75 were selected as candidate for passage-specific genes. Then, candidates of passage-specific genes for each passage were selected with criterion that the gene is highly expressed (z-score above zero) at this passage, and such high expression cannot be observed in more than two additional stages. These genes were selected as passage-specific genes.

To get a consistent landscape of chromatin accessibility, the merged accessible profile was created with mergeBed in bedtools v2.27.1 (Quinlan and Hall, 2010) and each region in the profile was annotated using ‘bedtools intersect’ with ‘-a’ being the region coordinate of merged profile and ‘-b’ being the peak coordinate of each passage collaborated with ‘-wa -wb’ to report the accessible region possession across all passages. The fragment coverage for all passages of accessible regions in the merged profile was computed using featureCounts v1.6.4 (Liao et al., 2014) with additional options “-F SAF -p -D 1000 --ignoreDup” and the count value was normalized with FPKM. Top 3,000 regions with highest variance in FPKM values across all passages were selected for PCA analysis. PCA analysis was carried out with prcomp function in R to compute PC1 and PC2 values of each passage.

### Single-cell RNA-sequencing and analysis

For single-cell RNA-seq (scRNA-seq) of K_D_L organoids, the organoids at passage 1, 3 and 8 were digested into single cell suspensions through gentle enzymatic dissociation with Tryple E for 5-10 min with at 37[. All single-cell suspensions were further prepared for sequencing according to 10x Chromium platform protocols found on: https://support.10xgenomics.com/single-cell-gene-expression.

10X Genomics sequencing data was processed using Cell Ranger Count (10X Genomics, v2.2.0 for organoid datasets and v3.1.0 for lung tumor datasets) to perform alignment with STAR v2.5.1b (Dobin et al., 2013), filtering, barcode counting, UMI counting and mapping UMIs to genes. Prebuilt mm10 reference genome v2.1.0 from 10X Genomics was used for read alignment and gene counting. Expression matrices of all samples were generated and merged with Seurat package v2.3.4 (Butler et al., 2018) followed by removing low-quality cells, respectively. Cells were excluded from further analysis based on the following criteria for organoid datasets: 1) the number of expressed genes lower than 2000 or larger than 7000; 2) the number of UMIs larger than 80,000; 3) 7.5% or more of UMIs were mapped to mitochondrial genes, and for tumor datasets: 1) the number of expressed genes lower than 300 or larger than 5000; 2) the number of UMIs lower than 500 or larger than 20,000; 3) 8% or more of UMIs were mapped to mitochondrial genes. Cells meeting the last threshold were usually non-viable or apoptotic. As a result, 16,492 genes in a total of 12,839 cells were detected for organoid datasets.

Data processing and visualization of the scRNA-seq expression matrices were performed with the Seurat package v2.3.4. The matrices were log normalized and scaled to 10,000 transcripts per cell by “NormalizeData” function. Uninteresting sources of cell-cell variation including the number of detected UMI as well as mitochondrial gene expression were removed by regression against each gene, and the resulting gene expression was scaled and centered by “ScaleData” function. Next, highly variable genes were identified by “FindVariableGenes” function with the minimum and maximum average gene expression cutoffs set to 0.0125 and 4 for organoid datasets and 0.1 and 4 for lung tumor datasets, and the minimum dispersion cutoff set to 0.5. Principal component analysis (PCA) was then carried out using the variable genes as input. 30 PCs were selected for organoid datasets for further analysis. Cell clusters were identified by the “FindClusters” function through a shared nearest neighbor modularity optimization, with cluster resolutions set to 0.8 and 12 cell clusters identified for organoid datasets. To perform dimensionality reduction analysis, the t-SNE and UMAP embeddings were calculated using “RunTSNE” and “RunUMAP” functions. Based on the normalized data, differentially expressed genes (DEGs) across cell clusters were identified by comparing expression values between cells of a given cluster and the remaining cells using “FindAllMarkers” function in Seurat based on the non-parametric Wilcox rank sum test with default parameter settings.

Pseudotime trajectory analysis was performed using the Monocle2 package v2.10.1(Trapnell et al., 2014). Briefly, Monocle2 was first used to estimate size factors and dispersions of genes. For organoid datasets, 1026 genes from bulk RNA-Seq with average expression within each passage increased or decreased monotonically along bulk organoid passages, which defined the order of organoid transition process, were used to order cells in pseudotime with Monocle2. For lung tumor datasets, we used DEGs from “FindAllMarkers” function in Seurat and filtered these genes with empirical dispersion above 0.1 to order cancer cells. Cells ordered along the trajectory were visualized in a reduced dimensional space with “plot_cell_trajectory” function in Monocle2. Differentially expressed genes along the pseudotime were calculated by “differentialGeneTest” function with q-value lower than 1e-50 and 1e-10 according to the distribution of log transformed q-value. Genes dynamically expressed along the pseudotime were clustered by “plot_pseudotime_heatmap” function with cluster number set to 8 for organoid datasets.

To calculate RNA velocity of single cells, the quantification of spliced and unspliced RNA were calculated from Cell Ranger derived BAM files using Python script velocyto.py v0.17.17 (La Manno et al., 2018), and then integrated into the Seurat object to generate input to downstream analysis while keeping cell filter and embedding consistent. RNA velocity analysis was performed using Scvelo package v0.2.2 (Bergen et al., 2020). Top 2000 highly variable genes were extracted and normalized followed by the first- and second-order moments calculation for velocity estimation with n_neighbors set to 40 and n_pcs set to 30 as above. The velocities were obtained by solving a stochastic model of transcriptional dynamics with “scv.tl.velocity” function. The velocity fields were projected onto the UMAP embedding using “scv.pl.velocity_embedding_stream” function. Cluster-grouped PAGA graph abstraction of velocity-inferred directionality on the UMAP embedding was generated with “scv.pl.paga” function.

Transcription factor activity in each cell was characterized by SCENIC v1.1.2.2 (Aibar et al., 2017) following the standard workflow with organoid datasets and cancer cells in lung tumor datasets. After an initial gene filter of the expression matrix, correlation networks between TFs and potential targets were calculated with GRNBoost2 in Arboreto package v0.1.5. Gene sets co-expressed with TFs were identified based on the correlation networks and compiled into modules (regulons). Next, cis-regulatory motif analysis were performed using RcisTarget with mm10 cisTarget databases v9 containing two gene-motif rankings: 10kb around the TSS (transcription start site), and 500bp upstream and 100bp downstream. Regulon activity was then scored using AUCell in each cell. Finally, the regulon activity score was projected onto low dimension space.

Individual cells were scored for the expression of gene signatures representing certain biological processes. The AUCell package v1.6.1 (Aibar et al., 2017) was used to calculate the enrichment score based on the AUC (Area Under the Curve) of the recovery curve for gene signatures in each cell.

### Patient specimen collection and transcriptomic analyses

A total of 68 patients out of 116 *KRAS^G12C^* lung cancer patients enrolled in KRYSTAL-1 trial (Jänne et al., 2022) had transcriptomic wide gene expression data available at the screening time point with outcome data and were included in the analysis. Three patients out of 116, and one patient out of 68 had SCC in pre-treatment biopsy. 14 and 17 of these 68 patients did not have *STK11* or *KEAP1* mutation status documented, respectively. The same criteria were used to call functional *STK11* and *KEAP1* mutations as in prior report (Jänne et al., 2022). Gene expression profiling was performed through HTG EdgeSeq platform (HTG Molecular Diagnostic, Inc.). Samples were profiled using the HTG Transcriptome Panel. Briefly, raw counts were normalized by library size and log2CPM expression values were calculated for each sample (edgeR v3.32.1). Genes with mean expression ≤ -1 were removed prior to signature analysis. SCC signature scores were calculated using GSVA (GSVA v1.38.2). GSVA was performed on log2CPM normalized data to generate signature scores including the Inamura Lung Cancer SCC Up and Down signatures (Inamura et al., 2005). Pearson correlation was used to measure the association of SCC signature scores, KRT6A expression and treatment duration. Survival curve analysis was done using survminer (v0.4.9) and survival (v3.1-12). The log-rank test was used to compare survival or treatment duration curves. All analyses were done in R (v4.0.2).

## Statistical analysis

For comparing means of 2 groups, two-tailed unpaired Student’s *t*-test with Welch’s correction was used. For comparing means of 3 groups, significance was determined using one-way ANOVA with Dunnett’s multiple comparisons test. Significance of grade of IHC staining was calculated by χ*2* test for trend. Student’s t-test and one-way ANOVA analyses were performed by Prism GraphPad software. χ*2* test was performed by R package (library (“coin”)). Differences with P < 0.05 were considered statistically significant. Data were all represented as mean ± SD or mean ± SEM.

## Data Availability Statement

The mouse lung cancer datasets have been deposited in the NODE database (project accessions OEP004095). The materials used in this study are available from the corresponding authors upon reasonable request.

## Illustration Tool

The graphical abstract image is created with BioRender.

## Acknowledgement

We thank Drs. T. Jacks, R. Depinho for providing the mice. We thank Drs. Daming Gao, Qibiao Wu, Gongwei Wu, Fei Li, Wangxin Guo, Wenming Liu, Juan He, Qiming Zhou, Jingyu Wang, Yuting Chen for helpful suggestions and technical supports.

