## Supplementary Figures for "Adeno-to-squamous transition drives resistance to KRAS inhibition in *LKB1* mutant lung cancer"

#### Figure S1

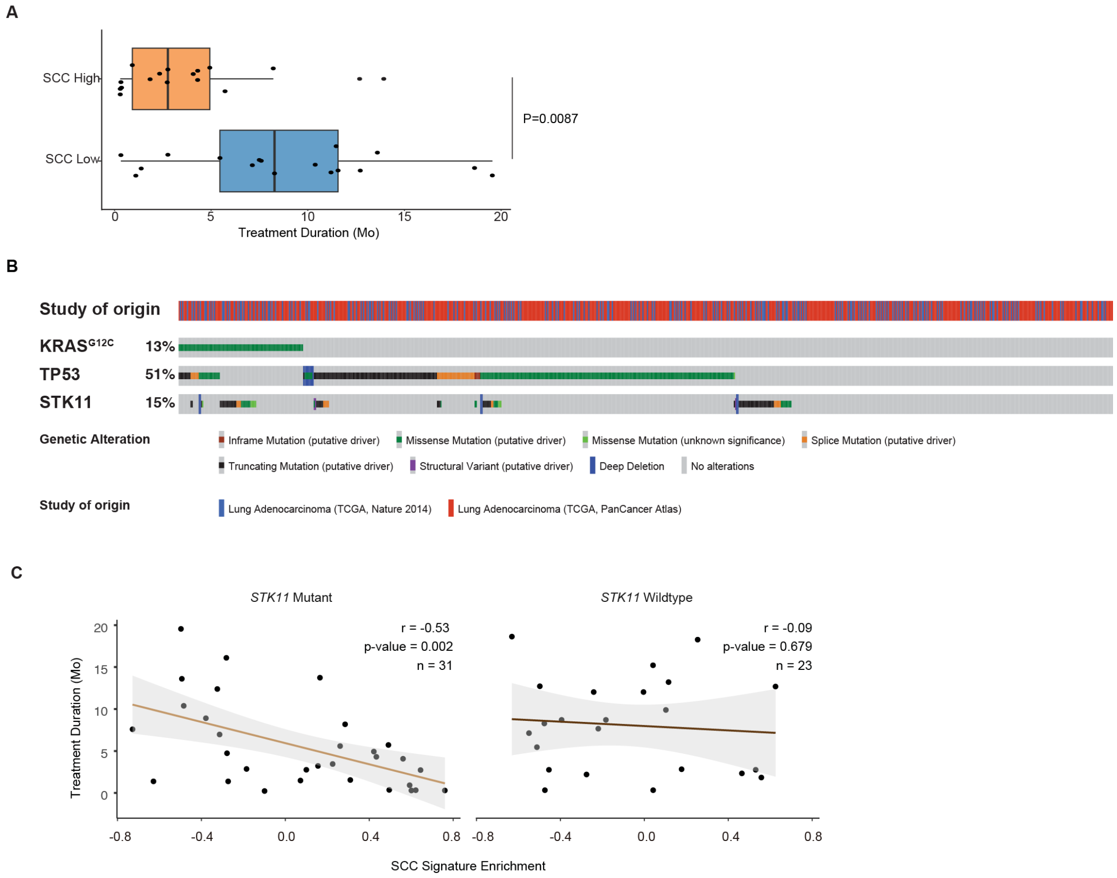


**Supplementary Figure S1. SCC signature correlates with poor clinical benefit from KRYSTAL1 clinical trial.**

**A.** Box plot showing range of treatment duration for patients stratified by SCC signature scores. High group included patients with in the upper quartile of SCC scores and low group included patients in the lower quartile of SCC scores. t-test was used for statistical analysis.

**B.** Somatic mutations of $KRAS^{G12C}$, *STK11/LKB1* and *TP53* in NSCLC patients from TCGA dataset.

**C.** Scatter plots showing the correlation between SCC signature and treatment duration for patients stratified by *STK11/LKB1* mutation status.

**Figure S2**

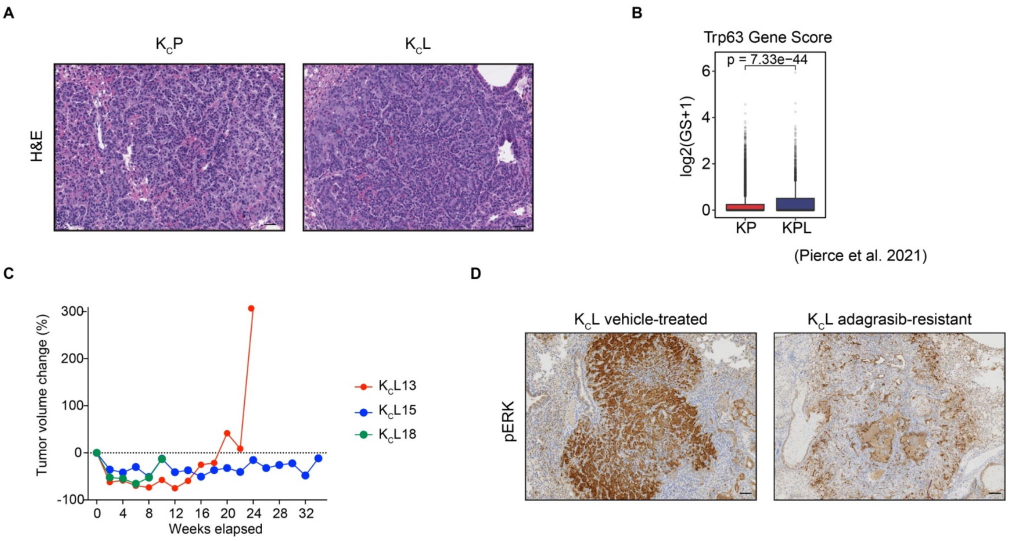


**Supplementary Figure S2. Establishment of adagrasib-resistant $K_C$P and $K_C$L mouse models.**

**A**. Representative photos of H&E staining in $K_C$P and $K_C$L mouse lung tumors. Scale bar, 50µm.

**B**. The gene scores of Trp63 in *Kras*/*Trp53* (KP) and *Kras*/*Trp53*/*Lkb1* (KPL) mouse lung tumors from scATAC-seq data.

**C.** Tumor volumes of $K_C$L mice monitored via MRI during adagrasib treatment. Tumor volume change is calculated as relative to tumor volume at the treatment initiation. Each line represents an individual mouse.

**D.** Representative photos of IHC staining in vehicle-treated and adagrasib-resistant $K_C$L tumors. Scale bar, 50µm.

**Figure S3**

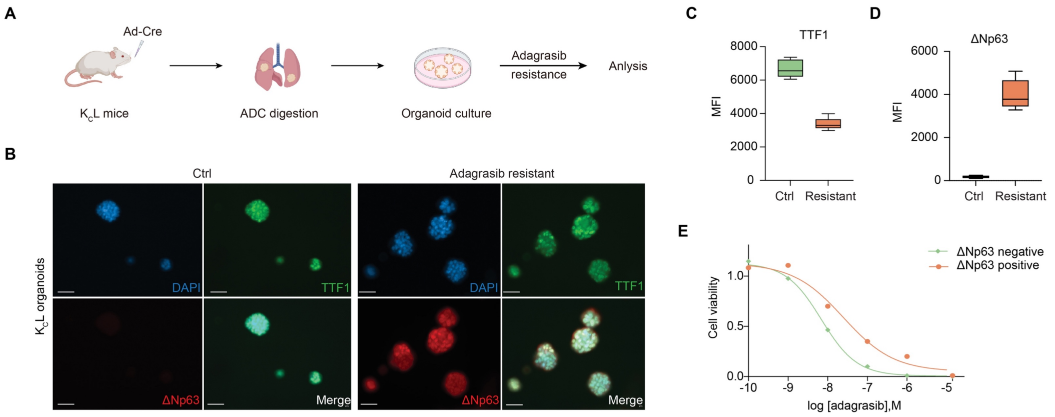


**Supplementary Figure 3. AST plasticity mediates resistance to KRAS inhibitor.**

**A.** Schematic illustration of experiments. Individual lung ADC from treatment-naïve $K_CL$ mice were used for the organoids culture with 12 weeks of adagrasib treatments (500nM).

**B-C.** Statistical analyses of the mean fluorescence intensity (MFI) of TTF1 (**B**) and ΔNp63 (**C**) in KCL organoids treated with Ctrl or adagrasib for 12 weeks (resistant). $p < 0.05$.

**D.** Representative photos of immunofluorescence staining for TTF1, ΔNp63 and DAPI in $K_CL$ organoids (Ctrl vs. Resistant). Scale bar, 50μm.

**E.** Cell viability of ΔNp63-negative and ΔNp63-positive organoids in response to adagrasib treatments.

#### Figure S4

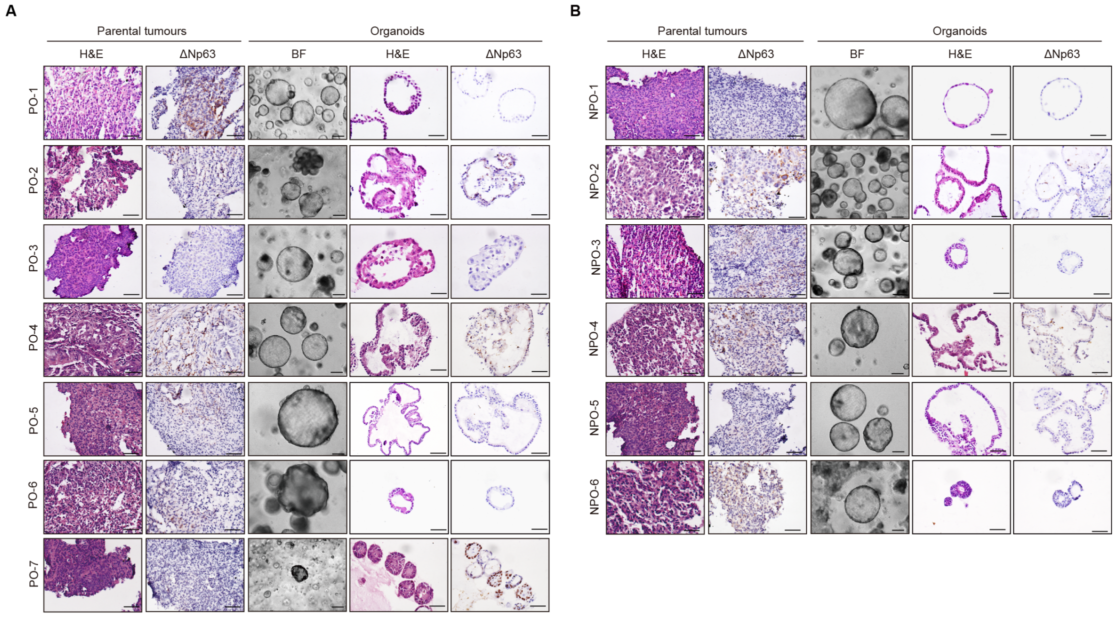


**Supplementary Figure S4. Establishment of mouse KDL lung ADC-derived organoids.**

**A.** Representative photos of H&E staining, ΔNp63 IHC staining, and/or bright field for KDL parental tumors (ADC) and plastic organoids (PO).

**B**. Representative photos of H&E staining, ΔNp63 immunostaining, and/or bright field for KDL parental tumors (ADC) and non-plastic organoids (NPO). Scale bar, 50μm.

**Figure S5**

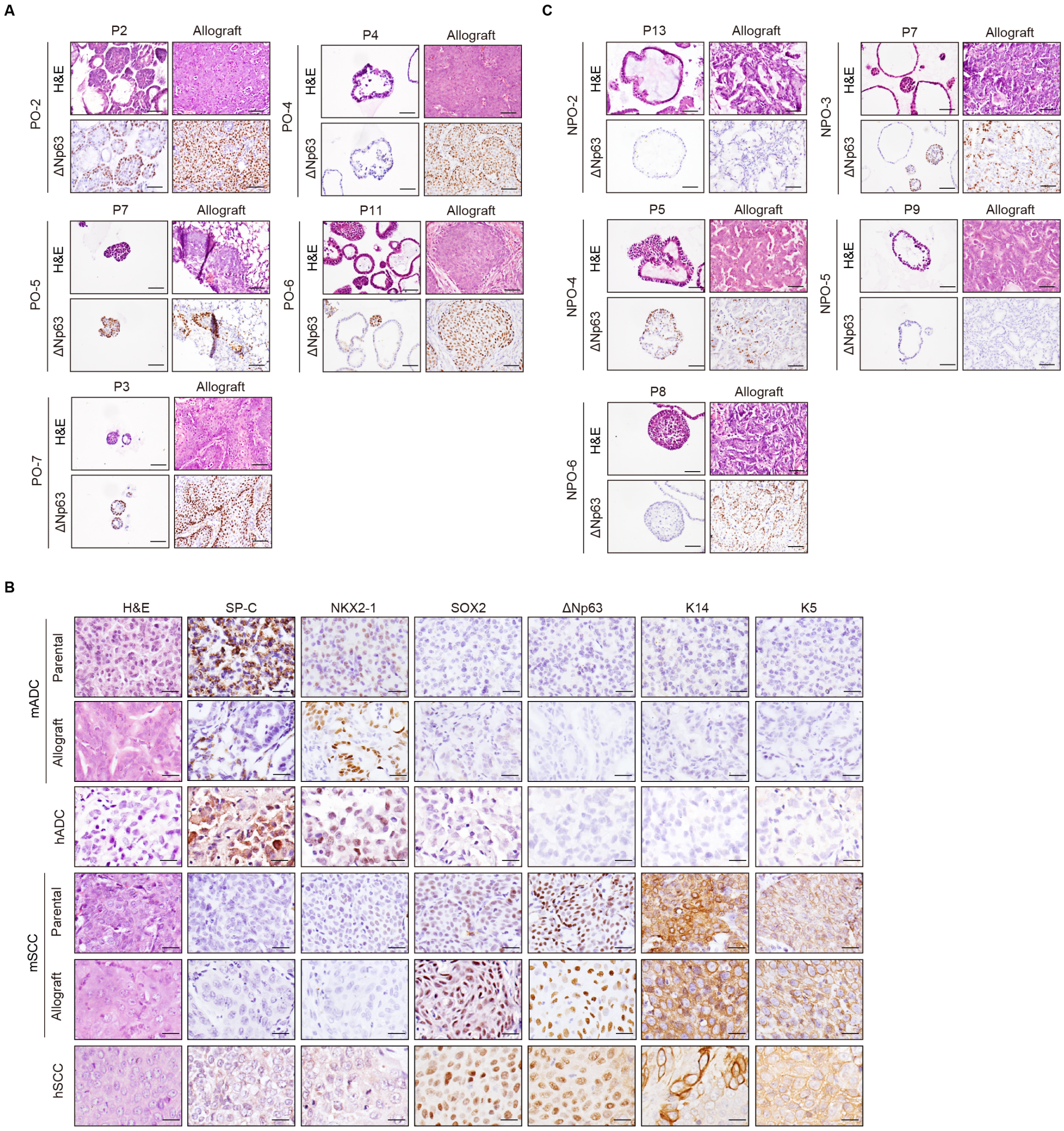


**Supplementary Figure S5. The organoids-derived allograft tumors closely recapitulate human tumors.**

**A**. Representative photos of H&E staining and ΔNp63 IHC staining for $K_DL$ plastic organoids at indicated passages and the corresponding allograft tumors. Scale bar, 50μm.

**B**. Representative photos of H&E and immunostaining for parental tumors (mouse ADC, mADC; mouse SCC, mSCC), mADC or mSCC organoids-derived allografts (allograft), human ADC (hADC), and human SCC (hSCC). Scale bar, 25μm.

**C**. Representative photos of H&E and immunostaining for non-plastic $K_DL$ ADC organoids at indicated passages and the corresponding allograft tumors. Scale bar, 50μm.

**Figure S6**

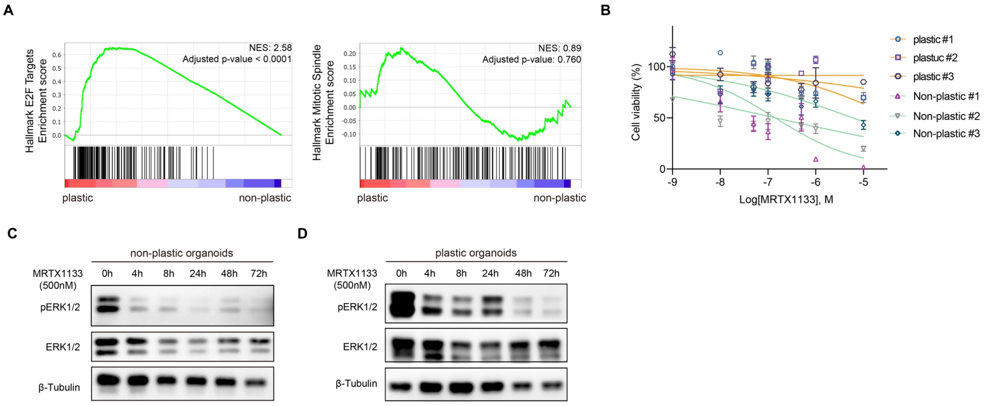


**Supplementary Figure S6. Comparative analyses of non-plastic and plastic $K_DL$ ADC organoids.**

**A**. GSEA analysis of proliferation-related signatures in non-plastic and plastic $K_DL$ ADC organoids.

**B**. Cell viability of plastic and non-plastic $K_DL$ ADC organoids treated with MRTX1133 (500nM).

**C-D.** Western blot for pERK1/2, ERK1/2 and GAPDH in non-plastic organoids (**C**) and plastic organoids (**D**) treated with MRTX1133.

**Figure S7**

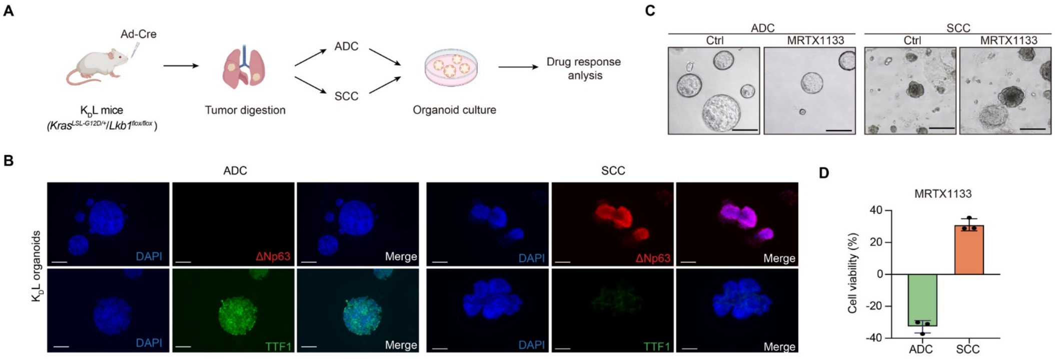


**Figure S7: Differential responses of $K_D$L ADC and SCC organoids to MRTX1133 treatments.**

**A.** Schematic illustration of experiments. We established organoids from the $K_D$L ADC and SCC and treated them with MRTX1133 at 500nM.

**B.** Representative photos of immunofluorescence staining for $K_D$L ADC and SCC organoids. Scale bar, 50µm.

**C.** Representative bright field photos of ADC and SCC organoids in ctrl group and MRTX1133-treated group.

**D.** Cell viability of ADC and SCC organoids when treated with MRTX1133 for 3 days.

**Figure S8**

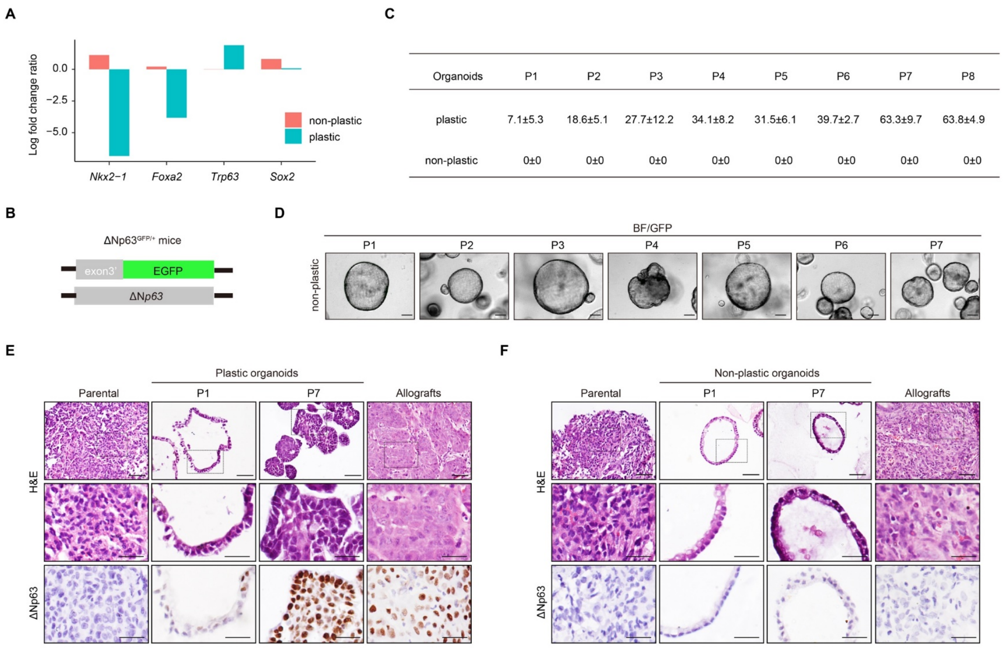


**Supplementary Figure S8. The establishment and characterization of the organoids system from *$Kras^{LSL\text{-}G12D/+}$;$Lkb1^{flox/flox}$;$\Delta Np63^{GFP/+}$* ($K_D$L-p63) mouse model.**

**A**. Relative expression of lineage-specific transcription factors in plastic and non-plastic organoids derived from mouse $K_D$L-p63 lung ADC.

**B**. Scheme illustrator of *$\Delta Np63^{GFP/+}$* mice. EGFP is knocked into exon 3 of *ΔNp63* allele and GFP positivity indicates the expression of *ΔNp63* gene.

**C**. Statistical analyses of $GFP^+$ organoids (%) in plastic organoids and non-plastic organoids with the passaging from P1 to P8.

**D.** Representative merged photos of bright field (BF) and fluorescence (GFP) of non-plastic organoids with the passaging from P1 to P7. Scale bar, 50μm.

**E.** Representative photos of H&E and ΔNp63 immunostaining for parental tumors, plastic organoids at different passages (P1, P7), and the corresponding allograft tumors. Scale bar, 50μm (top panel), 25μm (middle and bottom panels).

**F.** Representative photos of H&E and ΔNp63 immunostaining for parental tumors, non-plastic organoids at different passages (P1, P7), and the corresponding allograft tumors. Scale bar, 50μm (top panel), 25μm (middle and bottom panels).

#### Figure S9

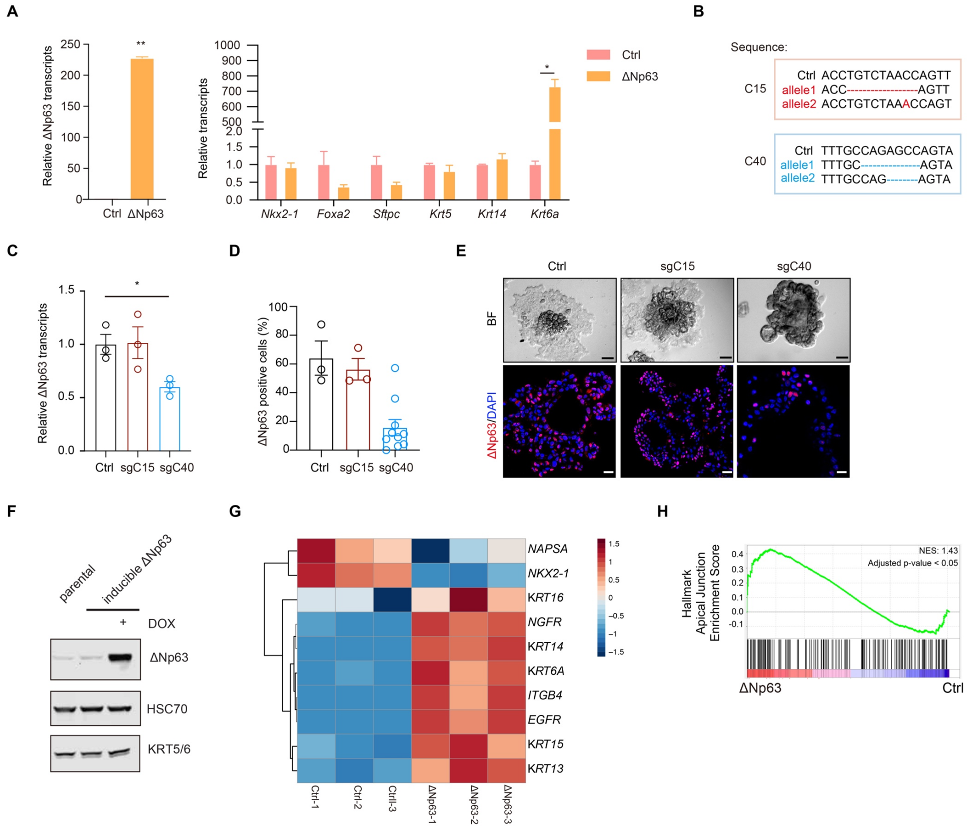


**Supplementary Figure S9. Molecular characterization of plastic $K_DL$ organoids and human KRAS$^{G12C}$-mutant cell line with modulation of ΔNp63 expression.**

**A**. Relative gene expression in mouse $K_DL$ ADC organoids after ΔNp63 overexpression.

**B**. The targeted gene sequences of C15 and C40 knockout.

**C.** Relative *ΔNp63* transcripts in plastic organoids transfected with sgC15 or sgC40.

**D**. Statistical analysis of ΔNp63-positive cells.

**E.** Representative photos of bright-field (BF) and ΔNp63 immunofluorescence staining for plastic organoids in ctrl, sgC15 and sgC40 groups. Scale bar, 50μm.

**F**. Western blot in human KRAS$^{G12C}$-mutant cell line H1373 with inducible ΔNp63 expression.

**G.** Heatmap showing the changes of ADC and SCC-related genes in H1373 cells with inducible ΔNp63 expression.

**H.** GSEA analysis of RNA-seq data for Hallmark Apical Junction pathway in H1373 cells with inducible ΔNp63 expression.

### Figure S10

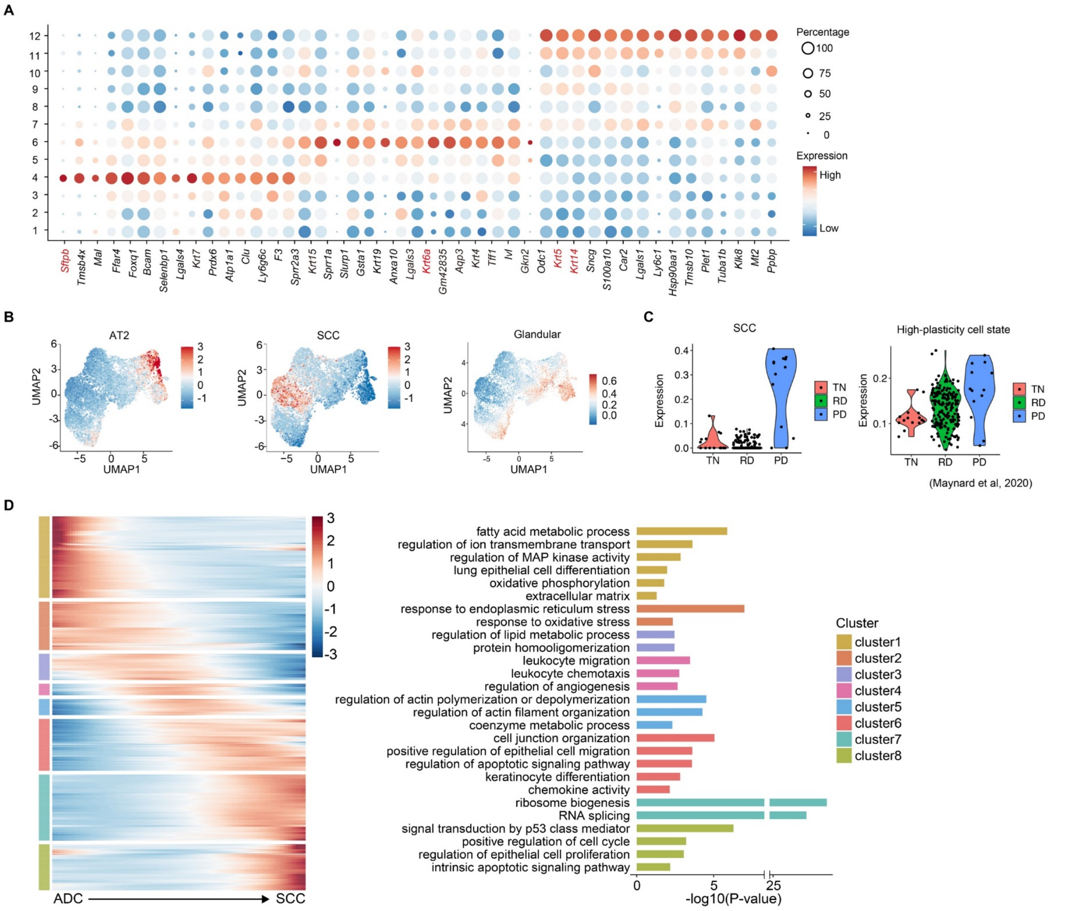


**Supplementary Figure S10. ScRNA-seq analysis of AST process in plastic K$_D$L organoids and human samples.**

**A.** Expression of highly enriched genes across different clusters in plastic K$_D$L organoids.

**B.** AUC scores of gene signatures of AT2, SCC and glandular signatures.

**C.** SCC signature and high-plasticity cell state signature enrichment in dynamic biopsy scRNA-seq data from a lung cancer patient with squamous transition from lung ADC. scRNA-Seq was performed with disease progression from treatment-naïve (TN), residual disease (RD) and progressive disease (PD).

**D.** GO pathway enrichment analyses of DEGs from scRNA-seq data of plastic K$_D$L organoids.

#### Figure S11

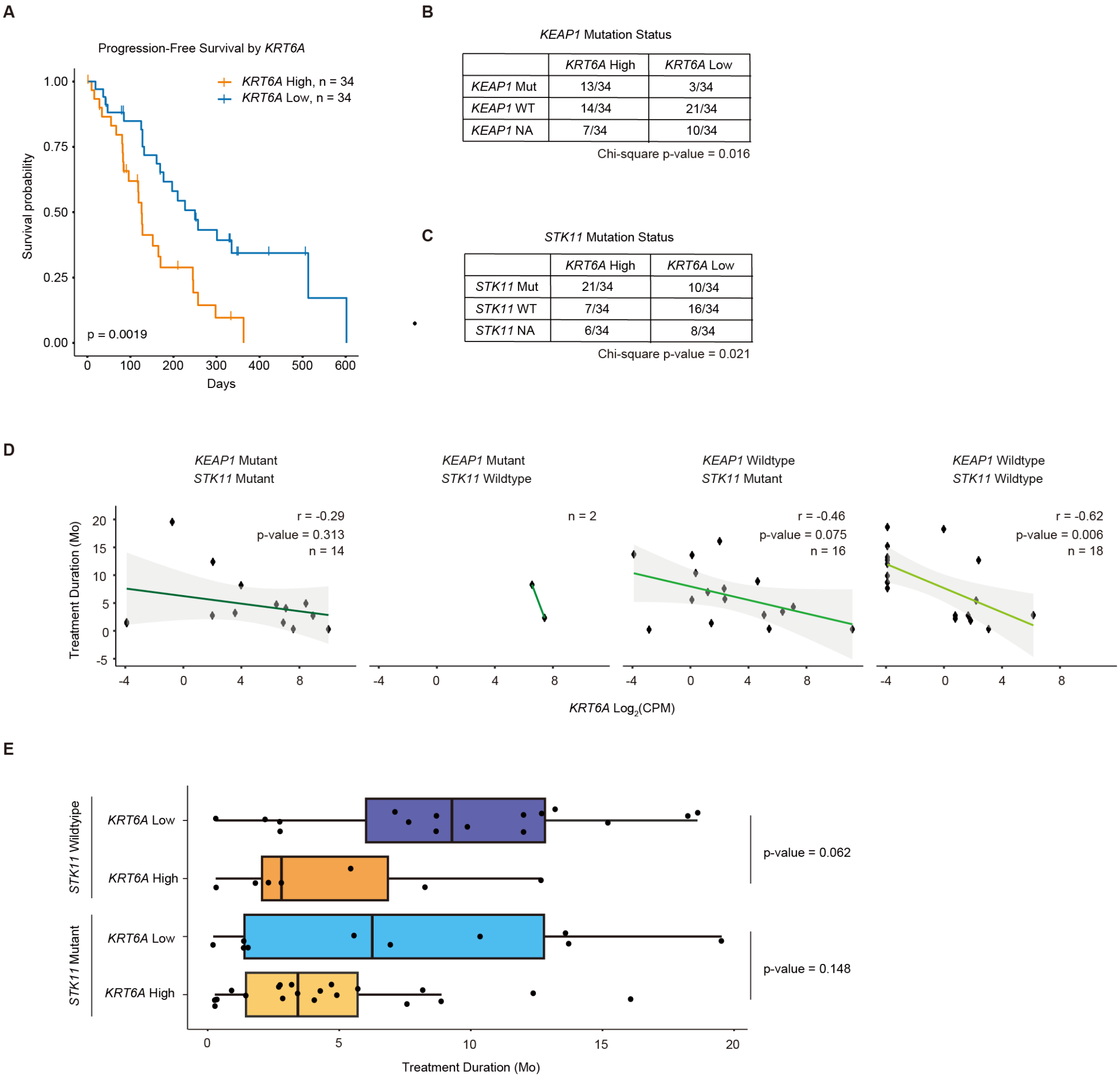


**Supplementary Figure S11. *KRT6A* expression correlation with clinical benefit from KRYSTAL1 clinical trial.**

**A.** Kaplan-Meier curve showing probability of survival for patients stratified by *KRT6A* expression above the median (high) and below the median (low). p-value = 0.0019.

**B-C**. Contingency table to classify patients with respect to subgroup of *KRT6A* expression and *KEAP1* (**B**) and *STK11* (**C**) mutation status.

**D.** Scatter plots showing correlation between *KRT6A* expression and treatment duration for patients stratified by *STK11* and *KEAP1* mutation status.

**E.** Boxplot showing range of treatment duration for patients stratified by *STK11* mutation status (mutant and wildtype) and *KRT6A* expression above median (high) and below the median (low). t-test p-values indicated on plot.

**Figure S12**

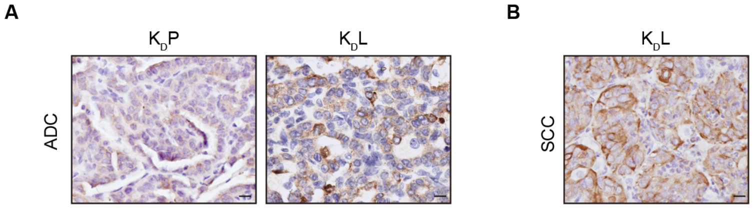


**Supplementary Figure S12. KRT6A expression in mouse tumors.**

**A-B.** Representative photos of KRT6A immunostaining in $K_D$P and $K_D$L ADC (**A**) and $K_D$L SCC (**B**). Scale bar, 50μm.
